# Single-Cell Mapping of tRNA Expression Dynamics Across Human Hematopoiesis

**DOI:** 10.64898/2026.08.14.744819

**Authors:** Marius Külp, Mireia Osuna Lopez, Anna-Sophia Wiegand, Sebastian P. Perner, Ronay Cetin, Yves Matthess, Stefan Günther, Tessa Schmachtel, Hendrik Schultheis, Mario Looso, Manuel Kaulich, Björn Häupl, Thomas Oellerich, Vladimir Benes, Halvard Bonig, Michael A. Rieger

## Abstract

How transfer RNA (tRNA) expression changes as hematopoietic stem cells are specified into human blood cell types remains unknown. tRNAs translate the 64-codon genetic code into the 21-amino acid protein code, and their relative abundance can influence proteomic output. Here, we introduce simultaneous single-cell tRNA and mRNA sequencing (sc-STM-seq), a scalable, high-throughput approach for profiling tRNAs alongside mRNA transcriptomes in individual cells. Applying sc-STM-seq to human bone marrow, we map tRNA expression and splicing across hematopoietic differentiation trajectories and identify tRNAs associated with stemness, differentiation, and specific cell lineages. Our study provides an atlas of the hematopoietic tRNA landscape and establishes tRNA expression as a previously underappreciated layer of cellular heterogeneity during human blood cell development.

## Introduction

Biological systems differ from inanimate physical systems in that, in addition to flows of energy and matter, they are governed by a flow of information(*1*): DNA makes RNA makes protein(*2*). Transfer RNAs (tRNAs) essentially support this flow of information by physically connecting RNA and protein. They decode triplets of RNA bases (codons) into amino acids and thus translate the 64-codon genetic code into the 21-amino acid protein code(*3*). The human genome contains 429 cytoplasmic tRNA genes, expressing more than 260 isotranscripts, representing 49 anticodon-sharing isodecoders, and 21 isoacceptor families transporting the same amino acid(*4*).

tRNA abundance was considered to be proportional to respective tRNA gene copy numbers, and thus largely constant across all cells of an organism(*5*).This notion became increasingly challenged by reports of differential tRNA expression across human cell types(*6–8*), and significant changes during differentiation of cancer cells(*9*), macrophages(*10*), and induced pluripotent stem cells(*11*). This variability is of biomedical relevance, as dysregulated tRNA expression has been associated with neurodevelopmental disorders(*12*) and cancer progression(*13*, *14*), i.e. through controlling translation of proliferation-regulating genes(*15*). Despite growing recognition of the importance of tRNA expression, large-scale resources such as the Human Cell Atlas still lack information on tRNA expression(*16*) as robust methods for single-cell tRNA profiling have not been available.

Hematopoiesis is the tightly regulated life-long regeneration of all blood cell types. Multipotent hematopoietic stem cells (HSCs) give rise to progressively committed progenitor populations, which differentiate into more than ten different blood cell lineages(*17*).

Recently, the importance of translational heterogeneity in both normal(*18*) and malignant(*19*) human hematopoiesis, mediated by a dynamic ribomethylome, has become increasingly apparent. Furthermore, matched single-cell proteomic and transcriptomic analyses have revealed quantitative differentiation-dependent discrepancies between protein and transcript abundances in the human hematopoietic stem and progenitor cell (HSPC) compartment(*20*), underscoring the contribution of translational regulation to hematopoietic cell fate.

Based on these observations, we reasoned that differential tRNA expression may represent an additional layer of translational regulation in the hematopoietic system. Therefore, we aimed to investigate whether tRNA expression changes dynamically during blood cell differentiation, and whether such changes occur in a lineage-specific manner. To address this, we developed simultaneous single-cell tRNA and mRNA sequencing (sc-STM-seq), a scalable, high-throughput approach for profiling cytosolic tRNAs and mRNAs in individual cells.

Applying sc-STM-seq to human bone marrow mononuclear cells (BMMNCs) we achieved robust quantification of full-length tRNAs alongside mRNAs, with strong concordance to bulk tRNA-sequencing. This enabled us to trace tRNA abundance along pseudotime-inferred differentiation trajectories from the HSC to the mature blood cell at single-cell resolution. We identify dynamic, lineage-associated tRNA expression patterns, uncover differential tRNA splicing, and define stemness- and differentiation-associated tRNAs.

Together, our study provides the first single-cell atlas of tRNA expression in human hematopoiesis and demonstrates that tRNA abundance and processing change dynamically during blood cell differentiation.

## Results

### sc-STM-seq simultaneously captures mRNAs and full-length tRNAs at single-cell resolution

tRNA-sequencing has been technically challenging due to rigid secondary structures, a high abundance of base modifications, and substantial sequence redundancy of tRNAs(*21*). Moreover, tRNAs lack poly(A) tails, rendering them inaccessible to many single-cell workflows. To overcome these limitations, we developed sc-STM-seq, a polyadenylation-independent method based on the iCELL8 single-cell system(*22*), in which individual cells serve as reaction compartments for reverse transcription and barcode incorporation (**Fig. 1A**).

**Fig. 1.**
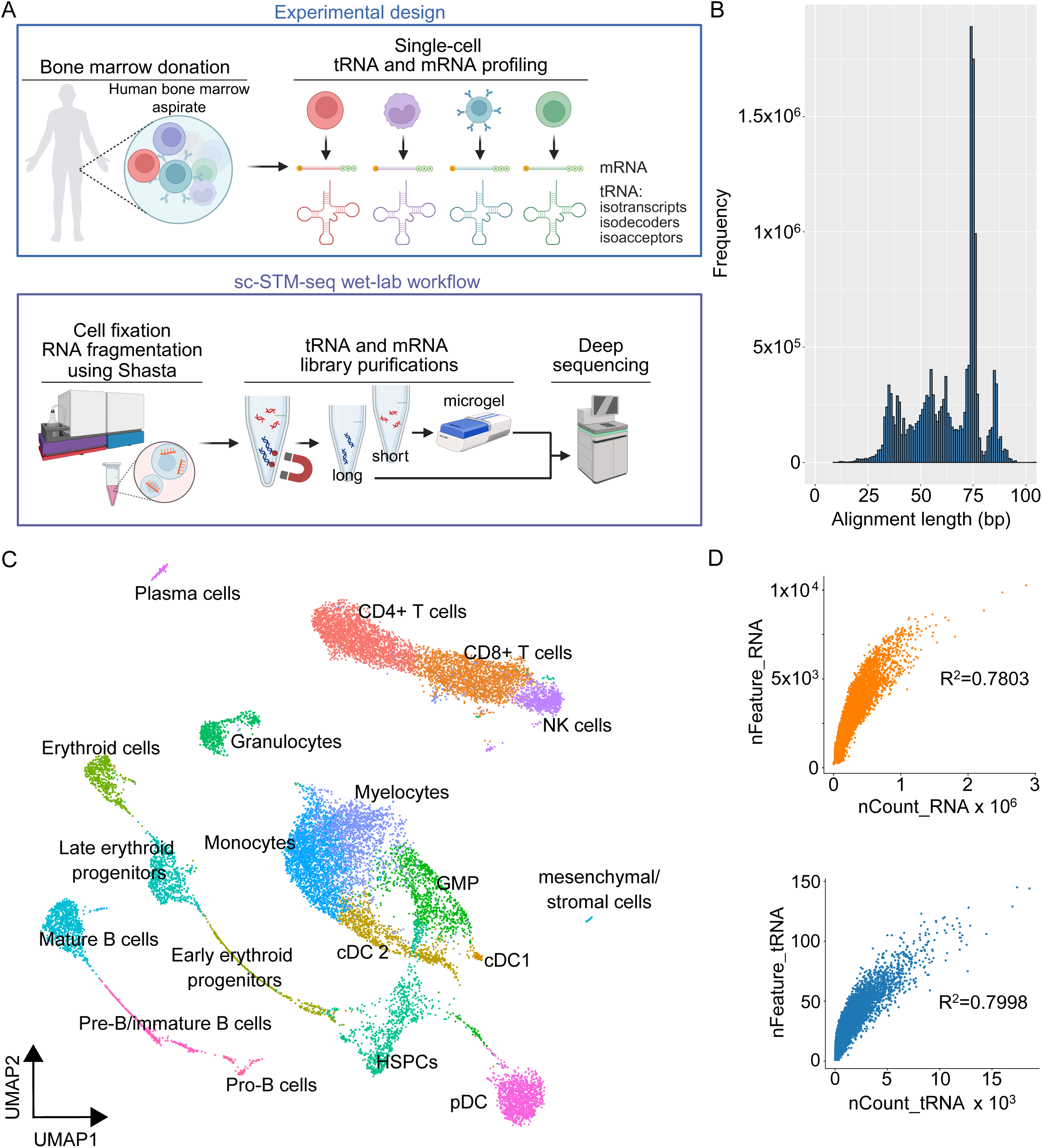
sc-STM-seq simultaneously captures mRNAs and full-length tRNAs at single-cell resolution. **A** Graphical abstract illustrating experimental design and sc-STM-seq wet-lab workflow. Created in BioRender. Rieger, A. (2026) https://BioRender.com/psd4nj8. **B** tRNA alignment histogram representing the number of tRNA reads against their alignment length. **C** UMAP projection with cell type annotation. HSPCs hematopoietic stem and progenitor cells. pDC plasmacytoid dendritic cells. GMP granulocyte-macrophage progenitors. **D** Feature/count correlations of the RNA assay (top) and the tRNA assay (bottom). R^2^ indicates Spearman correlation values.

Our approach distinguishes itself from the standard protocol by preserving tRNA-derived cDNAs during library purification while efficiently depleting primer dimers. In addition, we integrated our previously established tRNA alignment strategy(*23*) into a computational workflow for multi-omic single-cell data processing.

We applied sc-STM-seq to 21,702 BMMNCs obtained from a healthy male and a healthy female donor (**Tab. S1**). After quality filtering, 17,788 cells were retained for downstream analysis. Analysis of tRNA alignment length distribution showed a peak at 72 - 76 nt, and 75% of reads aligned to more than 50 nt, indicating efficient capture of full-length tRNAs, typically 72 - 85 nt in length (**Fig. 1B**).

Cells were clustered based on mRNA expression using principal component analysis (PCA) followed by UMAP projection, which identified 19 distinct clusters annotated by canonical hematopoietic marker genes (**Fig. 1C**). We captured a median of 149,773 mRNA reads per cell, corresponding to a median of 2,131 detected genes. All UMAP clusters were comprised of cells from both donors (**Fig. S1**). Cell numbers per cluster ranged from 123 (Pro-B cells) to 2,563 (CD4+ T cells) (**Tab. S2**). tRNAs were quantified at isotranscript, isodecoder (shared anticodon) and isoacceptor (shared amino acid) level. To allow the scientific community to access and visualize our tRNA expression atlas in detail, we established a web-based display publicly accessible at https://bioinformatics.mpi-bn.mpg.de/hematopoietic-tRNA-atlas.

To validate tRNA quantification, we compared our sc-STM-seq data from BMMNCs to peripheral blood (PB) bulk tRNA-seq results from ten healthy donors(*23*). Additionally, we performed bulk tRNA-seq on prospectively isolated CD3+CD4+ T cells and CD34+ HSPCs from mobilized PB (mPB) of three healthy donors, and compared these profiles to the corresponding single-cell clusters. In all cases, sc-STM-seq measurements showed strong concordance with bulk tRNA-seq data, with Spearman correlation coefficients between 0.81 and 0.84, p<0.0001 (**Fig. S2A**).

sc-STM-seq captured a median of 430 (1–17,392) tRNA reads per cell, representing a median of 16 (1–145) tRNA species (**Fig. S2B**). Highly, intermediately and lowly expressed tRNAs and mRNAs exhibited comparable read count distributions (**Fig. S3**).

Median tRNA reads per cell ranged from 53 in granulocytes to 1,948 in conventional dendritic cells (DCs) 1 (**Tab. S2**). Interestingly, DC subtypes displayed the highest tRNA contents, including conventional DC 1 and 2 with median values of 1,948 and 1,225.5 tRNAs/cell, respectively, and plasmacytoid DCs with 1,514 tRNAs/cell. HSPCs also contained high amounts of tRNAs (861). Overall, we observed a trend toward decreasing median tRNAs per cell during differentiation. Along the myeloid trajectory, median tRNA counts declined from HSPCs (861) to granulocyte-macrophage progenitors (GMPs, 540), myelocytes (311), and granulocytes (53). A similar pattern was observed during erythroid differentiation, with decreasing median tRNA counts from HSPCs (861) to early erythroid progenitors (679.5), late erythroid progenitors (262), and erythroid cells (236).

### Blood cell populations differ in tRNA expression and splicing

We performed differential tRNA expression analysis to compare tRNA profiles across distinct blood cell populations (**Fig. 2A**). This identified sets of differentially expressed tRNAs in multiple comparisons, including tRNA-Asp-GTC-3 in monocytes vs. CD8+ T cells, tRNA-Tyr-GTA-2 in B cells vs. erythroid cells, tRNA-Gln-TTG-4 in CD4+ vs. CD8+ T cells, and tRNA-Lys-TTT-4 in NK cells vs. CD8+ T cells, among others (**Fig. S4A**).

**Fig. 2.**
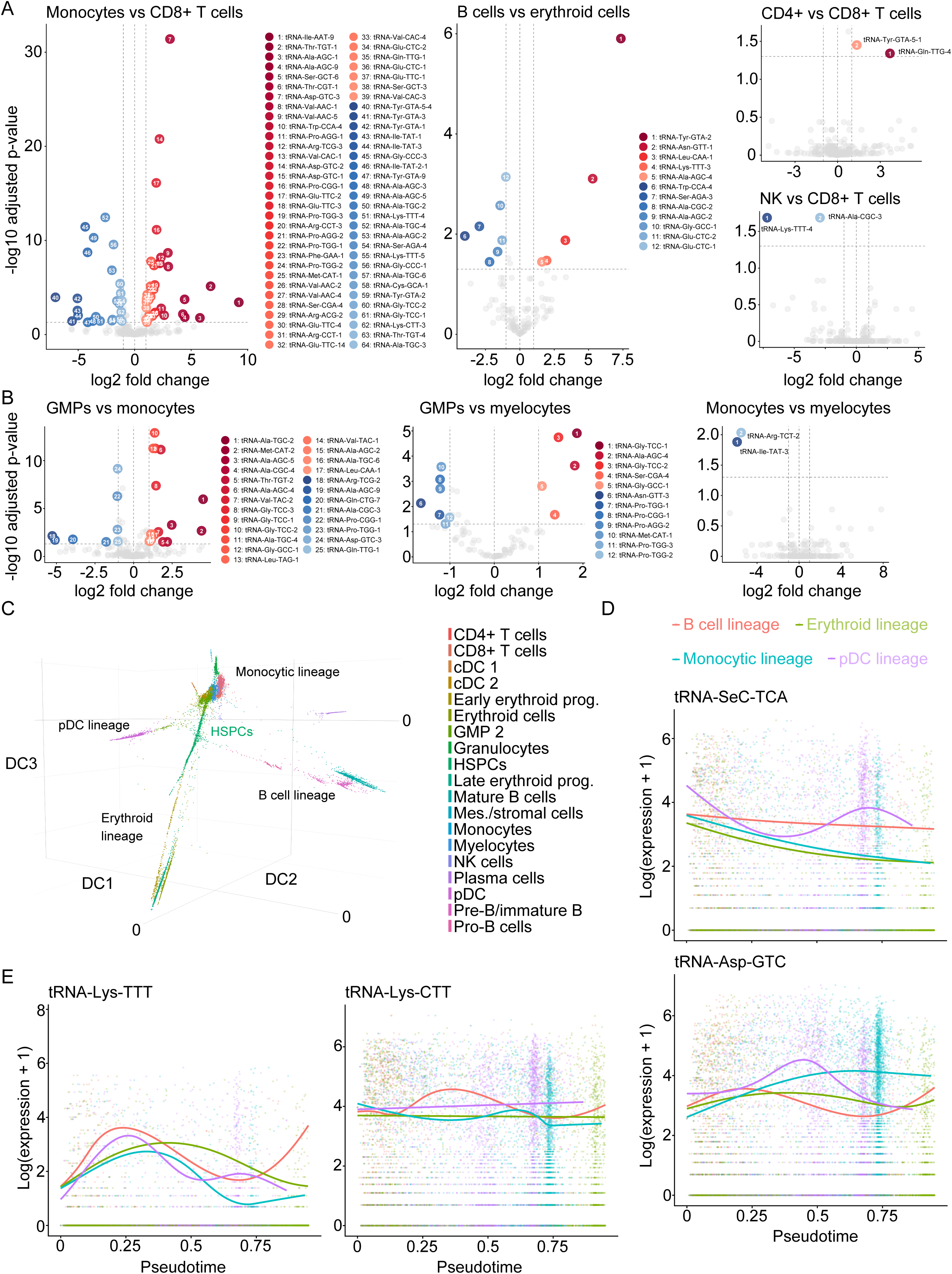
Blood cell populations differ in tRNA expression and splicing. **A** Volcano plots of differential tRNA expression comparisons of indicated cell types in pseudobulk. log2fc cut-off = 1, padj cut-off = 0.05. DEseq2 results are summarized in **supplementary file S1**. **B** Volcano plots of differential tRNA expression comparisons of GMP-descending cell types. DEseq2 results are summarized in **supplementary file S1**. **C** Diffusion map with cell type annotations and identified lineages. DC diffusion component. **D** Tradeseq analysis plots of tRNA-SeC-TCA and tRNA-Asp-GTC. **E** Tradeseq analysis plots of tRNA-Lys-TTT and tRNA-Lys-CTT.

Strikingly, closely related mature cell types, e.g. CD4+ and CD8+ T cells, displayed much fewer differentially expressed isotranscripts (only two tRNAs) compared to more distantly related populations from different lineages, such as monocytes and CD8+ T cells with 64 differentially expressed tRNAs (**Fig. 2A**). Moreover, the number of differentially expressed tRNAs decreased along the monocytic lineage, comparing GMPs with monocytes (25), GMPs with myelocytes (12), and monocytes with myelocytes (2) (**Fig. 2B**). These observations suggested that tRNA expression patterns may be linked to lineage identity and undergo dynamic changes during hematopoietic differentiation, analogous to well-characterized mRNA expression programs.

In addition to expression differences, we observed variation in tRNA processing. Notably, tRNA-Leu-CAA-4 exhibited differential splicing across cell populations (**Fig. S4B**), reflected by distinct ratios of intronic and exonic alignments among clusters (**Fig. S4C**). Additionally, alignment visualization suggested tolerance towards a misincorporation at position A58 which could potentially be introduced during reverse transcription of a modified nucleotide.

### tRNA expression changes dynamically along global differentiation trajectories

To investigate whether tRNA expression follows differentiation-dependent patterns along global trajectories, we constructed a diffusion map based on mRNA expression capturing continuous and discrete cell state transitions (**Fig. 2C**). This analysis resolved four major lineages: monocytic, erythroid, B cell and plasmacytoid DC (pDC). Lineage-specific diffusion maps enabled detailed assessment of mRNA and tRNA dynamics along pseudotime (**Fig. S5**).

We validated the inferred trajectories by observing progressive downregulation of HSC marker genes, including *CRHBP* and *PROM1*, during differentiation (**Fig. S6A**). As expected, lineage-specific marker genes increased along pseudotime: *EBF1* in the B cell lineage, *TFRC* encoding CD71 in erythroid cells, *ITGAM* encoding CD11b in the monocytic lineage, and *CLEC4C* in pDCs (**Fig. S6B**).

Using pseudotime trajectory analysis, we observed pronounced dynamics in tRNA isodecoder expression along differentiation paths (**Fig. S6C**). Notably, tRNA-SeC-TCA (**Fig. 2D**) displayed a stemness-associated pattern, characterized by progressive downregulation along monocytic and erythroid trajectories, and showed a transient decrease followed by late re-expression during pDC differentiation, while exhibiting modest decline in the B cell lineage. In contrast, tRNA-Asp-GTC showed a continuous increase along the monocytic trajectory until reaching saturation, while displaying only transient upregulation in other lineages. tRNA-Lys-TTT (**Fig. 2E**) exhibited early transient upregulation followed by mid-pseudotime downregulation across all lineages. Notably, its isoacceptor, tRNA-Lys-CTT, remained relatively stable but was expressed at substantially higher levels.

A dot plot of isodecoder expression across cell types confirmed enrichment of tRNA-SeC-TCA in HSPCs and strong expression of tRNA-Asp-GTC in myeloid cell types (**Fig. S7**). Similar patterns were observed at isoacceptor level for selenocysteine and aspartate tRNAs (**Fig. S8**).

As the importance of selenoproteins for HSC function has been demonstrated in a tRNA-SeC knockout mouse model(*24*), we sought to assess selenoprotein abundance in human HSPCs. Quantitative proteomics of human mPB CD34+ HSPCs in comparison to CD4+ T cells revealed enrichment of the selenocysteine-specific elongation factor EEFSEC and glutathione peroxidase 1 (GPX1) in HSPCs, whereas mitochondrial glutathione reductase (MT-GSR) and selenophosphate synthetase 2 (SEPHS2) were more abundant in T cells (**Fig. S9**).

In summary, these analyses identified stem cell-associated expression of tRNA-SeC-TCA and characterized tRNA-Asp-GTC as strongly associated with monocytic differentiation. The increased abundance of tRNA-SeC-TCA and EEFSEC suggested an increased capacity for selenoprotein biosynthesis in HSPCs.

### tRNA expression changes during early HSPC differentiation

We observed a distinct tRNA expression profile in HSPCs compared to all other hematopoietic populations, despite the considerable heterogeneity within the HSPC compartment (**Fig. 3A**). To investigate whether tRNA expression changes during early hematopoietic differentiation, we re-clustered the HSPC subset and resolved highly immature cell states, including HSCs and multiple progenitor populations, consistent with our previous study(*25*) (**Fig. 3B**). This enabled detailed comparisons between closely related HSPC subtypes.

**Fig. 3.**
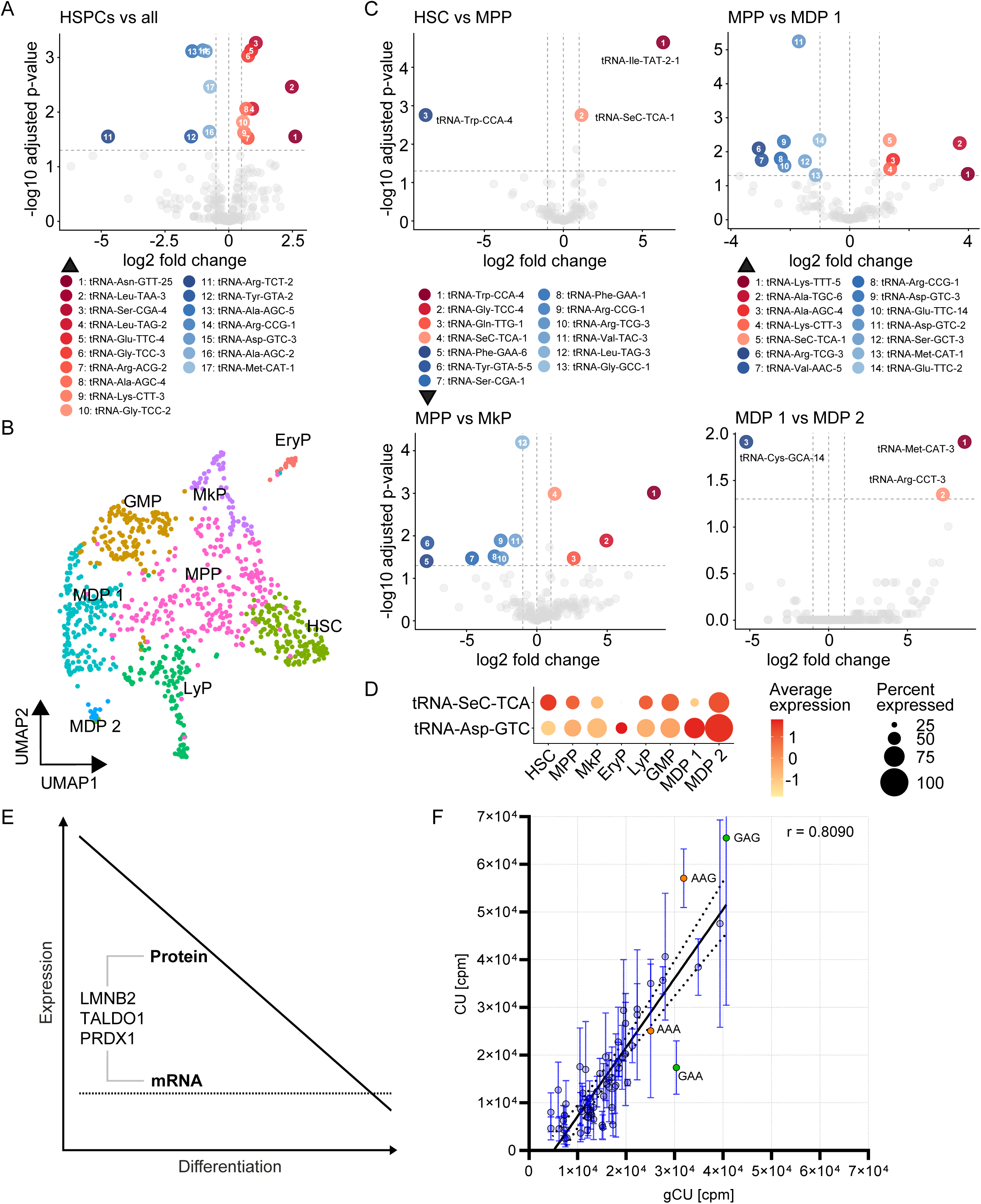
tRNA expression changes during early HSPC differentiation. **A** Volcano plot indicating differentially expressed tRNAs comparing HSPCs with all other cell types. log2fc cut-off = 0.5, padj cut-off = 0.05. DEseq2 results are summarized in **supplementary file S2**. **B** UMAP projection of re-clustered HSPCs with cell annotations. HSC hematopoietic stem cell. MPP multipotent progenitor. EryP erythroid progenitor. MkP megakaryocytic progenitor. GMP granulocyte-macrophage progenitor. MDP monocytic-dendritic progenitor. LyP Lymphoid progenitor. **C** Volcano plots indicating differentially expressed tRNAs of HSPC subpopulations. log2fc cut-off = 1, padj cut-off = 0.05. DEseq2 results are summarized in **supplementary file S2**. **D** Dotplot visualizing isodecoder expression and percentage of expressing cells of tRNA-SeC-TCA and tRNA-Asp-GTC across HSPC subpopulations. **E** Schematic representation of quantitative protein-transcript discrepancies of LMNB2, TALDO1, and PRDX1 during GMDP differentiation as described by Furtwängler *et al.*(*20*). **F** Codon usage (CU) analysis of LMNB2, TALDO1 and PRDX1 open-reading-frames showed AAG enrichment and GAA depletion compared to the human genomic codon usage (gCU). Lys codons are depicted in orange, Glu codons highlighted in green. Spearman correlation r=0.8090. Dots indicate the mean value per codon and bars show the standard error of mean (S.E.M.). The line represents a simple linear regression with 95% confidence interval indicated by the dotted lines. CU codon usage. gCU genomic codon usage. cpm codons per million.

Strikingly, tRNA expression changed during the earliest differentiation steps from HSCs to multipotent progenitors (MPPs), including reduced expression of tRNA-SeC-TCA-1, further supporting its stemness-association (**Fig. 3C**). tRNA-SeC-TCA-1 expression further declined during differentiation from MPPs toward megakaryocytic progenitors (MkPs) and monocytic-dendritic progenitors (MDPs). In contrast, the abundance of tRNA-Asp-GTC-2 and tRNA-Asp-GTC-3 increased during MPP-to-MDP differentiation, underpinning the association of tRNA-Asp-GTC isodecoders with myeloid differentiation. The stemness-associated expression pattern of tRNA-SeC-TCA and the myeloid differentiation-associated expression of tRNA-Asp-GTC were consistently observed across the entire HSPC hierarchy (**Fig. 3D**). Together, these analyses demonstrate that the tRNA expression dynamics observed along global hematopoietic trajectories are already established during the earliest HSPC differentiation stages.

### tRNA expression and codon usage may explain quantitative protein-transcript discrepancies during HSPC differentiation

Furtwängler and colleagues recently identified a GMDP differentiation-associated decline in LMNB2, TALDO1, and PRDX1 protein abundance without corresponding changes in transcript levels, using matched single-cell proteomics and transcriptomics of human HSPCs(*20*) (**Fig. 3E**).

We hypothesized that differential tRNA expression may contribute to this protein-transcript discrepancy and therefore analyzed the codon usage of the respective open-reading-frames. Compared with human genomic codon usage, these transcripts showed relative enrichment of the lysine codon AAG and relative depletion of the glutamate codon GAA (**Fig. 3F** and **S10A**). Importantly, during MPP-to-MDP differentiation, expression of the AAG-decoding tRNA-Lys-CTT-3 decreased, whereas the GAA-decoding tRNAs tRNA-Glu-TTC-2 and −14 increased (**Fig. 3C**). These data suggest that tRNA abundance may constrain AAG-dependent translation in MDPs and GAA-dependent translation in MPPs, resulting in higher translation efficiency of AAG-enriched and GAA-depleted open-reading-frames in MPPs, followed by reduced protein output during differentiation toward MDPs.

Consistent with this model, we found increased TALDO1 and PRDX1 protein abundances in HSPCs compared to T cells using quantitative proteomic analysis (**Fig. S10B**). This was accompanied by elevated expression of the AAG-encoding tRNAs tRNA-Lys-CTT-2, −3, and −5 as quantified by bulk tRNA-sequencing (**Fig. S10C**). These findings suggest that translation of TALDO1 and PRDX1 may likewise be constrained by tRNA availability during lymphoid differentiation.

Together, our observations identify AAG as a differentiation-associated translational bottleneck and GAA as a stemness-associated translational bottleneck, providing a potential explanation for declining LMNB2, TALDO1, and PRDX1 protein abundances despite stable transcript levels during GMDP differentiation.

### Reduced tRNA abundance alters the proteome in a codon-specific manner

To investigate how reduced tRNA abundance influences the proteome, we knocked down the glutamate and aspartate tRNAs tRNA-Asp-GTC-1, −2, tRNA-Glu-TTC-2, tRNA-Glu-CTC-1, and −2 in the pro-B cell line SEM using CRISPR/Cas9 in comparison to a non-targeting control (NTC). tRNA-seq confirmed knockdown efficiencies ranging from 27% to 53%, with few off-target effects and a compensatory upregulation of lysine isodecoders (**Fig. S11A**, **supplementary file S5**). To assess proteome-wide consequences, we performed quantitative proteomics under steady-state conditions (DMSO) and following 1h of proteasome inhibition (MG132) to capture proteins undergoing enhanced degradation (**Fig. S11B**). Under steady-state conditions, 327 proteins were upregulated and 536 downregulated, whereas MG132 treatment revealed 210 upregulated and 502 downregulated proteins (**Fig. S11C, D**, **supplementary file S6**). Most differentially abundant proteins changed concordantly in both conditions (**Fig. S11E**).

We next asked whether these proteomic changes reflected altered codon demand. Codon usage analysis revealed that downregulated proteins were enriched for GAG (Glu) and AAG (Lys) codons, whereas upregulated proteins were depleted of GAT (Asp) and AAA (Lys) codons; both groups were depleted of GAA (Glu) codons (**Fig. S12A,B**). Consistent with these codon-level changes, amino acid composition analysis showed that upregulated proteins contained fewer aspartate residues, while both up- and downregulated proteins were depleted of glutamate residues compared with unchanged proteins (**Fig. S12C,D, supplementary file S6**). Thus, proteins affected by reduced availability of glutamate and aspartate tRNAs were enriched or depleted of the respective codons and depleted of corresponding amino acids.

These findings suggest that experimentally reducing tRNA abundance reshapes the proteome according to the codon composition of translated transcripts, indicating tRNA availability to be a determinant of proteomic output.

### tRNA profiles indicate distinct cell types

To benchmark sc-STM-seq against bulk tRNA-seq (**Fig. S2A**), we sorted CD34+CD38low HSPCs, CD34+CD38high progenitors and CD3+CD4+ T cells from mPB of three healthy donors (**Fig. S13A**) and analyzed their tRNA profiles in bulk. PCA based solely on tRNAs clearly separated HSPCs and progenitors from T cells, a result that was corroborated by unsupervised clustering analysis (**Fig. S13B**). These findings implied that tRNA expression profiles may contribute to cell identity and could be used to distinguish cell types.

To inspect this at single-cell resolution, we performed normalization, scaling, clustering, and UMAP projection of sc-STM-seq data using tRNA isotranscript, isodecoder and isoacceptor expression (**Fig. 4A**). This analysis identified 14 - 17 clusters. Mapping previously annotated cell types onto these tRNA-based clusters revealed non-random distributions across all levels of tRNA aggregation. HSPCs were strongly enriched within a single dominant cluster, whereas CD4+ T cells were more broadly distributed without a predominant cluster association (**Fig. 4B**). Interestingly, GMPs displayed broad distribution across multiple clusters at the isotranscript level, but became more restricted at the isodecoder and isoacceptor levels, similar to HSPCs (**Fig. 4C**). In contrast, monocytes and CD4+ T cells exhibited relatively even distributions across clusters at all aggregation levels and were least represented within the HSPC-enriched cluster.

**Fig. 4.**
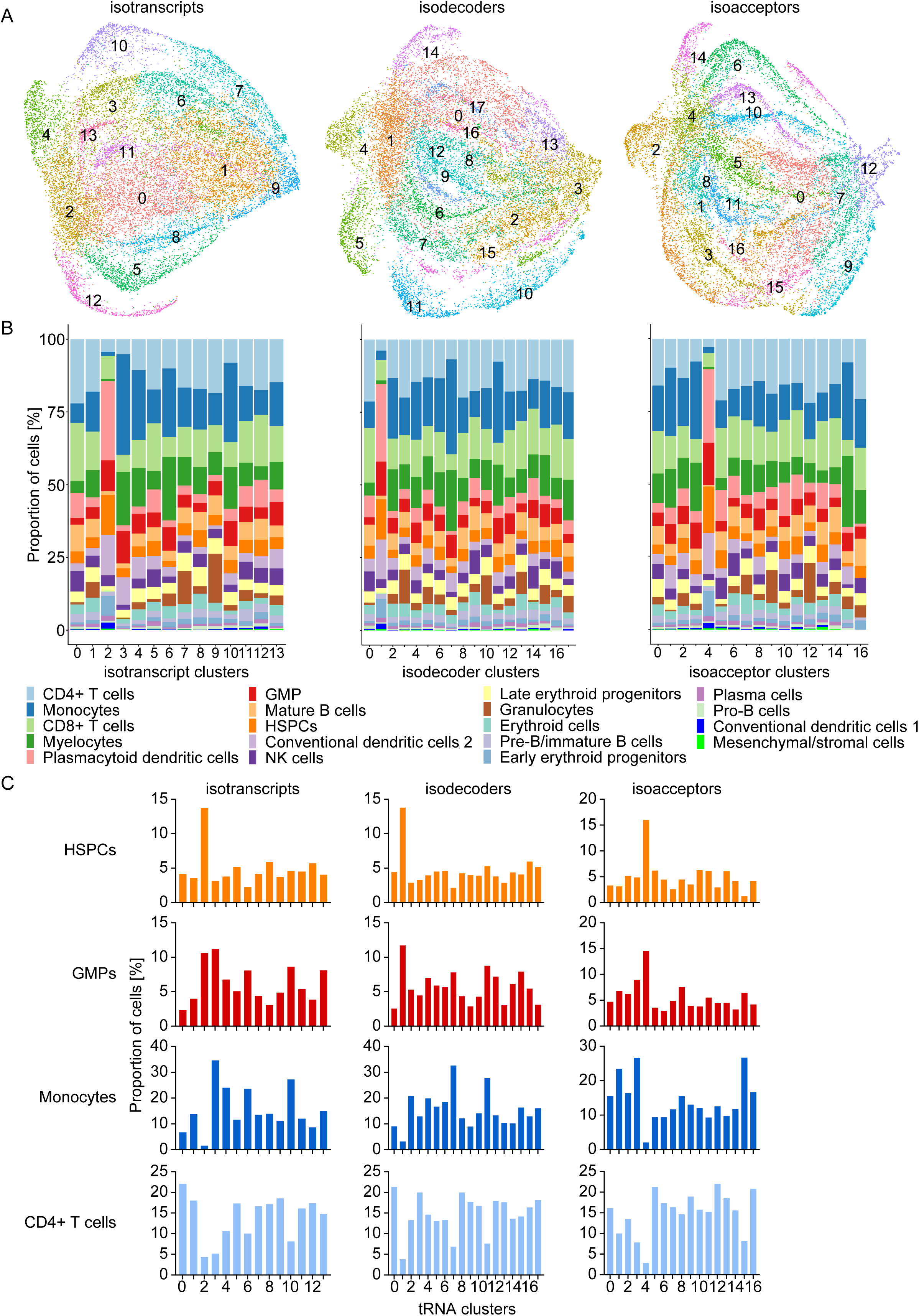
tRNA profiles indicate distinct cell types. **A** UMAPs based on tRNA expression at isotranscript, isodecoder, and isoacceptor level. **B** Bar plots indicate the proportion of cell types belonging to each tRNA-based cluster. **C** Bar plots indicating the proportion of HSPCs, GMPs, monocytes, and CD4+ T cells in tRNA-based clusters.

Accordingly, these results indicated that tRNA expression contributed to cell identity in a cell type-dependent manner. While tRNA profiles were highly distinctive for certain populations, such as HSPCs, they were less discriminative for others, including T cells and monocytes.

## Discussion

The genetic code is degenerate, with multiple synonymous codons encoding the same amino acid(*26*). Importantly, codon-specific tRNA availability directly affects translation elongation rates, as ribosomes must pause longer when cognate tRNAs are scarce(*27–31*). This establishes a regulatory role for codon usage, as codon composition affects mRNA stability, translational fidelity, and protein folding, leading to the concept of codon usage as a “secondary genetic code”(*32*) the rules of which, however, we currently do not fully understand.

With sc-STM-seq, we introduce a high-throughput approach to interrogate this secondary genetic code at single-cell resolution across entire cellular systems such as human hematopoiesis. Our method robustly captures full-length tRNAs while minimizing co-detection of tRNA-derived fragments, which have distinct biological functions(*33*).

While previous single-cell methods for non-coding RNA-sequencing reported incidental capture of tRNAs(*34–36*), they did not demonstrate robust recovery of full-length tRNAs, systematic benchmarking against bulk tRNA-sequencing, or comprehensive analysis across hematopoietic differentiation trajectories. Moreover, none of these studies included human bone marrow or HSPCs.

In addition to quantifying tRNA abundance, sc-STM-seq enables the detection of differential tRNA splicing and exhibits tolerance to misincorporations, likely arising from base modifications. For example, the observed misincorporation in tRNA-Leu-CAA-4 occurred at a known methylation site (m1A58, according to GtRNAdb(*4*)). However, further methodological development and validation will be required before sc-STM-seq can be reliably used to map tRNA modifications(*37*).

We identify tRNA-SeC-TCA as a stemness-associated tRNA, with highest expression in immature HSPCs and progressive downregulation during differentiation. Strikingly, the murine counterpart of this tRNA has been shown to be a critical regulator of HSC function, where its knockout impairs hematopoiesis, alters lineage commitment, and reduces self-renewal capacity by abolishing selenoprotein synthesis(*24*). Additionally, our observation of increased abundance of the selenocysteine-specific elongation factor EEFSEC in HSPCs underscores the importance of selenoprotein biosynthesis for HSPCs and suggests their enhanced capacity to translate selenocysteine. Together, these findings point to a conserved and biologically meaningful role for tRNA-SeC-TCA not only in murine but also human hematopoietic regulation.

Our study suggests that the tRNAome contributes to translational heterogeneity of the hematopoietic system, representing an additional regulatory layer alongside the dynamic ribomethylome(*18*, *19*). This finding is important, as our data indicate that the interplay between tRNA abundance and codon usage may explain quantitative discrepancies between protein and transcript abundances during hematopoietic differentiation(*20*). Beyond affecting protein abundance and co-translational folding(*38*), dynamic changes in tRNA availability may also cause amino acid misincorporations (*39*).

Finally, we previously reported that tRNA supply is highly conserved across healthy individuals(*23*), suggesting the applicability of our findings despite the limited number of donors analyzed here.

In summary, sc-STM-seq provides a scalable platform for single-cell tRNA profiling and enables the systematic investigation of tRNA biology across complex cellular systems. Our findings reveal that tRNA abundance and processing are dynamically regulated during hematopoietic differentiation. These results establish tRNAs as a previously underappreciated layer of cellular heterogeneity and identity in human blood cell development.

## Acknowledgments

We thank Martine Pape and Marion Bodach of DKTK Proteomics Platform Frankfurt and Uwe Plessmann of Bioanalytical Mass Spectrometry Group, MPI-NAT Göttingen (Prof. Henning Urlaub), for technical assistance with sample preparation and LC/MS measurements. We thank Anjali Cremer and Sarah Bröchtel, Goethe University Hospital Frankfurt, for kindly providing Cas9-expressing SEM cells.

## Funding

European Hematology Association KOG-202409-06466 (MK)

Deutsche Jose Carreras Leukämie-Stiftung DJCLS15 R/2023 (MAR)

Deutsche Forschungsgemeinschaft DFG RI 2462/9-1 and RI 2462/10-1 (MAR)

LOEWE Hessian Funding Program Hessen State Ministry for Higher Education, Research and the Arts, III L5 − 519/03/03.001 – [0015] and III 5.7 - 519/03/10.001-(0004) (MAR, HB)

Deutsche Forschungsgemeinschaft DFG ExStra EXC2026 Translational Hub 2 (ML)

Deutsche Forschungsgemeinschaft DFG under Germanýs Excellence Strategy – EXC 2026, Cardio-Pulmonary Institute, Project ID: 390649896. (MAR, MK, MaK)

## Author contributions

Conceptualization: MK, MAR

Methodology: MK, MOL, AW, SPP, RC, YM, SG, TS, HS, ML, MKa, BH, TO, VB, HB, MAR

Investigation: MK, MOL, AW, SPP, RC, YM, SG, TS, HS, ML, MKa, BH, TO, VB, HB, MAR

Visualization: MK Software: MK, HS, ML

Funding acquisition: MK, MAR

Project administration: MK, MAR

Supervision: MAR

Resources: HB

Writing – original draft: MK, MAR

Writing – review & editing: MK, MOL, AW, SPP, RC, YM, SG, TS, HS, ML, MKa, BH, TO, VB, HB, MAR

## Supplementary Materials

Materials and Methods

Supplementary Files S1 to S6

Supplementary Tables S1 and S2

Figs. S1 to S13

## References and Notes

1. K. L. Hoke, S. L. Zimmer, A. B. Roddy, M. J. Ondrechen, C. E. Williamson, N. R. Buan, Reintegrating Biology Through the Nexus of Energy, Information, and Matter. Integr Comp Biol 61, 2082–2094 (2021).

2. F. Crick, Central Dogma of Molecular Biology. Nature 227, 561–563 (1970).

3. H. F. Noller, The ribosome comes to life. Cell 187, 6486–6500 (2024).

4. P. P. Chan, T. M. Lowe, GtRNAdb 2.0: an expanded database of transfer RNA genes identified in complete and draft genomes. Nucleic Acids Research 44, D184–D189 (2016).

5. P. M. Sharp, W. H. Li, The codon Adaptation Index--a measure of directional synonymous codon usage bias, and its potential applications. Nucleic Acids Res 15, 1281–1295 (1987).

6. O. Pinkard, S. McFarland, T. Sweet, J. Coller, Quantitative tRNA-sequencing uncovers metazoan tissue-specific tRNA regulation. Nat Commun 11, 4104 (2020).

7. K. A. Dittmar, J. M. Goodenbour, T. Pan, Tissue-Specific Differences in Human Transfer RNA Expression. PLOS Genetics 2, e221 (2006).

8. C. Scheepbouwer, E. Aparicio-Puerta, C. Gomez-Martin, H. Verschueren, M. van Eijndhoven, L. E. Wedekind, S. Giannoukakos, N. Hijmering, L. Gasparotto, H. T. van der Galien, R. S. van Rijn, E. Aronica, R. Kibbelaar, V. M. Heine, P. Wesseling, D. P. Noske, W. P. Vandertop, D. de Jong, D. M. Pegtel, M. Hackenberg, T. Wurdinger, A. Gerber, D. Koppers-Lalic, ALL-tRNAseq enables robust tRNA profiling in tissue samples. Genes Dev 37, 243–257 (2023).

9. H. Gingold, D. Tehler, N. R. Christoffersen, M. M. Nielsen, F. Asmar, S. M. Kooistra, N. S. Christophersen, L. L. Christensen, M. Borre, K. D. Sørensen, L. D. Andersen, C. L. Andersen, E. Hulleman, T. Wurdinger, E. Ralfkiær, K. Helin, K. Grønbæk, T. Ørntoft, S. M. Waszak, O. Dahan, J. S. Pedersen, A. H. Lund, Y. Pilpel, A Dual Program for Translation Regulation in Cellular Proliferation and Differentiation. Cell 158, 1281–1292 (2014).

10. K. Van Bortle, D. H. Phanstiel, M. P. Snyder, Topological organization and dynamic regulation of human tRNA genes during macrophage differentiation. Genome Biol 18, 180 (2017).

11. L. Gao, A. Behrens, G. Rodschinka, S. Forcelloni, S. Wani, K. Strasser, D. D. Nedialkova, Selective gene expression maintains human tRNA anticodon pools during differentiation. Nat Cell Biol 26, 100–112 (2024).

12. R. Ishimura, G. Nagy, I. Dotu, H. Zhou, X.-L. Yang, P. Schimmel, S. Senju, Y. Nishimura, J. H. Chuang, S. L. Ackerman, Ribosome stalling induced by mutation of a CNS-specific tRNA causes neurodegeneration. Science 345, 455–459 (2014).

13. H. Goodarzi, X. Liu, H. C. B. Nguyen, S. Zhang, L. Fish, S. F. Tavazoie, Endogenous tRNA-Derived Fragments Suppress Breast Cancer Progression via YBX1 Displacement. Cell 161, 790–802 (2015).

14. Z. Zhang, Y. Ye, J. Gong, H. Ruan, C.-J. Liu, Y. Xiang, C. Cai, A.-Y. Guo, J. Ling, L. Diao, J. N. Weinstein, L. Han, Global analysis of tRNA and translation factor expression reveals a dynamic landscape of translational regulation in human cancers. Commun Biol 1, 234 (2018).

15. L. B. Earnest-Noble, D. Hsu, S. Chen, H. Asgharian, M. Nandan, M. C. Passarelli, H. Goodarzi, S. F. Tavazoie, Two isoleucyl tRNAs that decode synonymous codons divergently regulate breast cancer metastatic growth by controlling translation of proliferation-regulating genes. Nat Cancer 3, 1484–1497 (2022).

16. J. E. Rood, S. Wynne, L. Robson, A. Hupalowska, J. Randell, S. A. Teichmann, A. Regev, The Human Cell Atlas from a cell census to a unified foundation model. Nature 637, 1065–1071 (2025).

17. M. A. Rieger, T. Schroeder, Hematopoiesis. Cold Spring Harb Perspect Biol 4, a008250 (2012).

18. O. Rabany, S. Ben Dror, M. Arafat, H. Aharoni Levitanus, Y. Halperin, V. Marchand, N. Romanovski, N. Ussishkin, M. Livneh Golany, A. Reches, J. Wexler, N. Mayorek, G. Monderer-Rothkoff, S. Shifman, W. M. Bouhou, M. VanInsberghe, C. Pauli, C. Müller-Tidow, O. Karmi, Y. Livneh, A. van Oudenaarden, Y. Motorin, D. Nachmani, Dynamic rRNA methylation regulates translation in the hematopoietic system and is essential for stem cell fitness. Blood 147, 520–533 (2026).

19. F. Zhou, N. Aroua, Y. Liu, C. Rohde, J. Cheng, A.-K. Wirth, D. Fijalkowska, S. Göllner, M. Lotze, H. Yun, X. Yu, C. Pabst, T. Sauer, T. Oellerich, H. Serve, C. Röllig, M. Bornhäuser, C. Thiede, C. Baldus, M. Frye, S. Raffel, J. Krijgsveld, I. Jeremias, R. Beckmann, A. Trumpp, C. Müller-Tidow, A Dynamic rRNA Ribomethylome Drives Stemness in Acute Myeloid Leukemia. Cancer Discov 13, 332–347 (2023).

20. B. Furtwängler, N. Üresin, S. Richter, M. B. Schuster, D. Barmpouri, H. Holze, A. Wenzel, K. Grønbæk, K. Theilgaard-Mönch, F. J. Theis, E. M. Schoof, B. T. Porse, Mapping early human blood cell differentiation using single-cell proteomics and transcriptomics. Science 390, eadr8785 (2025).

21. N. H. Padhiar, U. Katneni, A. A. Komar, Y. Motorin, C. Kimchi-Sarfaty, Advances in methods for tRNA sequencing and quantification. Trends Genet 40, 276–290 (2024).

22. L. Liu, X. Dong, Y. Tu, G. Miao, Z. Zhang, L. Zhang, Z. Wei, D. Yu, X. Qiu, Methods and platforms for analysis of nucleic acids from single-cell based on microfluidics. Microfluid Nanofluidics 25, 87 (2021).

23. M. Külp, H. Bonig, M. A. Rieger, Variation of Human Transfer RNA Demand and Supply. bioRxiv [Preprint] (2026). 10.64898/2026.02.26.708130.

24. Y. Aoyama, H. Yamazaki, K. Nishimura, M. Nomura, T. Shigehiro, T. Suzuki, W. Zang, Y. Tatara, H. Ito, Y. Hayashi, Y. Koike, M. Fukumoto, A. Tanaka, Y. Zhang, W. Saika, C. Hasegawa, S. Kasai, Y. Kong, Y. Minakuchi, K. Itoh, M. Yamamoto, S. Toyokuni, A. Toyoda, T. Ikawa, A. Takaori-Kondo, D. Inoue, Selenoprotein-mediated redox regulation shapes the cell fate of HSCs and mature lineages. Blood 145, 1149–1163 (2025).

25. H. Komic, T. Schmachtel, C. Simoes, M. Külp, W. Yu, A. Jolly, M. S. Nilsson, C. Gonzalez, F. Prosper, H. Bonig, B. Paiva, F. B. Thorén, M. A. Rieger, Continuous map of early hematopoietic stem cell differentiation across human lifetime. Nat Commun 16, 2287 (2025).

26. M. Nirenberg, Historical review: Deciphering the genetic code – a personal account. Trends in Biochemical Sciences 29, 46–54 (2004).

27. C.-H. Yu, Y. Dang, Z. Zhou, C. Wu, F. Zhao, M. S. Sachs, Y. Liu, Codon Usage Influences the Local Rate of Translation Elongation to Regulate Co-translational Protein Folding. Mol Cell 59, 744–754 (2015).

28. A. Dana, T. Tuller, The effect of tRNA levels on decoding times of mRNA codons. Nucleic Acids Research 42, 9171–9181 (2014).

29. D. Chu, D. J. Barnes, T. von der Haar, The role of tRNA and ribosome competition in coupling the expression of different mRNAs in Saccharomyces cerevisiae. Nucleic Acids Res 39, 6705–6714 (2011).

30. K. S. Koutmou, A. Radhakrishnan, R. Green, Synthesis at the Speed of Codons. Trends in Biochemical Sciences 40, 717–718 (2015).

31. D. Chu, E. Kazana, N. Bellanger, T. Singh, M. F. Tuite, T. von der Haar, Translation elongation can control translation initiation on eukaryotic mRNAs. EMBO J 33, 21–34 (2014).

32. G. Hanson, J. Coller, Codon optimality, bias and usage in translation and mRNA decay. Nat Rev Mol Cell Biol 19, 20–30 (2018).

33. B. Kuhle, Q. Chen, P. Schimmel, tRNA renovatio: Rebirth through fragmentation. Mol Cell 83, 3953–3971 (2023).

34. A. Isakova, D. D. Liu, I. Cvijović, R. Sinha, A. E. Eastman, S. Saul, A. M. Detweiler, N. Neff, S. Einav, I. L. Weissman, S. R. Quake, Scalable single-cell total RNA sequencing unifies coding and noncoding transcriptomics. Nat Biotechnol, 1–12 (2026).

35. F. Salmen, J. De Jonghe, T. S. Kaminski, A. Alemany, G. E. Parada, J. Verity-Legg, A. Yanagida, T. N. Kohler, N. Battich, F. van den Brekel, A. L. Ellermann, A. M. Arias, J. Nichols, M. Hemberg, F. Hollfelder, A. van Oudenaarden, High-throughput total RNA sequencing in single cells using VASA-seq. Nat Biotechnol 40, 1780–1793 (2022).

36. F. B. Dinçaslan, S. W. Y. Ngang, R. Z. Tan, L. F. Cheow, Automated high-throughput profiling of single-cell total transcriptome with scComplete-seq. Nucleic Acids Res 53, gkaf699 (2025).

37. A. Behrens, G. Rodschinka, D. D. Nedialkova, High-resolution quantitative profiling of tRNA abundance and modification status in eukaryotes by mim-tRNAseq. Molecular Cell 81, 1802–1815.e7 (2021).

38. S. Pechmann, J. Frydman, Evolutionary conservation of codon optimality reveals hidden signatures of cotranslational folding. Nat Struct Mol Biol 20, 237–243 (2013).

39. S. Tsour, R. Machné, A. Leduc, S. Widmer, E. Koo, J. Guez, K. J. Karczewski, N. Slavov, Alternate RNA decoding results in stable and abundant proteins in mammals. Nature, 1–10 (2026).

40. S. Bräuninger, K. Thorausch, B. Luxembourg, M. Schulz, K. U. Chow, E. Seifried, H. Bonig, Deferrals of volunteer stem cell donors referred for evaluation for matched-unrelated stem cell donation. Bone Marrow Transplant 49, 1419–1425 (2014).

41. T. Schmachtel, H. Bonig, M. A. Rieger, FACS-Based Assessment of Human Hematopoietic Stem and Progenitor Cells. Int J Mol Sci 26, 8381 (2025).

42. T. Metsalu, J. Vilo, ClustVis: a web tool for visualizing clustering of multivariate data using Principal Component Analysis and heatmap. Nucleic Acids Res 43, W566–570 (2015).

43. M. Martin, Cutadapt removes adapter sequences from high-throughput sequencing reads. EMBnet.journal 17, 10–12 (2011).

44. The Galaxy Community, The Galaxy platform for accessible, reproducible, and collaborative data analyses: 2024 update. Nucleic Acids Res 52, W83–W94 (2024).

45. STAR: ultrafast universal RNA-seq aligner | Bioinformatics | Oxford Academic. https://academic.oup.com/bioinformatics/article/29/1/15/272537?login=false.

46. Y. Liao, G. K. Smyth, W. Shi, featureCounts: an efficient general purpose program for assigning sequence reads to genomic features. Bioinformatics 30, 923–930 (2014).

47. B. Langmead, C. Trapnell, M. Pop, S. L. Salzberg, Ultrafast and memory-efficient alignment of short DNA sequences to the human genome. Genome Biol 10, R25 (2009).

48. B. Langmead, S. L. Salzberg, Fast gapped-read alignment with Bowtie 2. Nat Methods 9, 357–359 (2012).

49. Y. Hao, S. Hao, E. Andersen-Nissen, W. M. Mauck, S. Zheng, A. Butler, M. J. Lee, A. J. Wilk, C. Darby, M. Zager, P. Hoffman, M. Stoeckius, E. Papalexi, E. P. Mimitou, J. Jain, A. Srivastava, T. Stuart, L. M. Fleming, B. Yeung, A. J. Rogers, J. M. McElrath, C. A. Blish, R. Gottardo, P. Smibert, R. Satija, Integrated analysis of multimodal single-cell data. Cell 184, 3573–3587.e29 (2021).

50. Single-cell proteo-genomic reference maps of the hematopoietic system enable the purification and massive profiling of precisely defined cell states | Nature Immunology. https://www.nature.com/articles/s41590-021-01059-0.

51. P. Angerer, L. Haghverdi, M. Büttner, F. J. Theis, C. Marr, F. Buettner, destiny: diffusion maps for large-scale single-cell data in R. Bioinformatics 32, 1241–1243 (2016).

52. L. Haghverdi, M. Büttner, F. A. Wolf, F. Buettner, F. J. Theis, Diffusion pseudotime robustly reconstructs lineage branching. Nat Methods 13, 845–848 (2016).

53. K. Van den Berge, H. Roux de Bézieux, K. Street, W. Saelens, R. Cannoodt, Y. Saeys, S. Dudoit, L. Clement, Trajectory-based differential expression analysis for single-cell sequencing data. Nat Commun 11, 1201 (2020).

54. P. Danecek, J. K. Bonfield, J. Liddle, J. Marshall, V. Ohan, M. O. Pollard, A. Whitwham, T. Keane, S. A. McCarthy, R. M. Davies, H. Li, Twelve years of SAMtools and BCFtools. Gigascience 10, giab008 (2021).

55. J. T. Robinson, H. Thorvaldsdóttir, W. Winckler, M. Guttman, E. S. Lander, G. Getz, J. P. Mesirov, Integrative genomics viewer. Nat Biotechnol 29, 24–26 (2011).

56. M. I. Love, W. Huber, S. Anders, Moderated estimation of fold change and dispersion for RNA-seq data with DESeq2. Genome Biol 15, 550 (2014).

57. H. Schultheis, J. Detleffsen, R. Wiegandt, M. Bentsen, Y. Alayoubi, G. Valente, M. F. Keßler, B. Bruns, D. Mirza, A. Usanayo, J. Walter, P. Goymann, M. Hobein, C. Kuenne, M. Looso, SC-Framework: A robust and FAIR semi-interactive environment for single-cell resolution datasets. bioRxiv [Preprint] (2025). 10.1101/2025.11.11.687874.

58. I. Virshup, S. Rybakov, F. J. Theis, P. Angerer, F. A. Wolf, anndata: Access and store annotated data matrices. Journal of Open Source Software 9, 4371 (2024).

59. K. Li, Z. Ouyang, Y. Chen, J. Gagnon, D. Lin, M. Mingueneau, W. Chen, D. Sexton, B. Zhang, Cellxgene VIP unleashes full power of interactive visualization and integrative analysis of scRNA-seq, spatial transcriptomics, and multiome data. bioRxiv [Preprint] (2022). 10.1101/2020.08.28.270652.

60. J. C. Enssle, B. Häupl, A. Qoku, B. Wang, G. W. Wright, S. Barrans, Y. Zhou, M. A. Care, C. Burton, C. Gribbin, J. Ziello, J. Weirather, Y. Dai, A. Kizhakeyil, X. Li, J. D. Phelan, S. Kanangat, S. Eckert, S. Scheich, S. Wolf, D. W. Huang, J. Jakob, S. P. Perner, A. Di Fonzo, M. Pape, M. Bodach, D. Jahn, U. Plessmann, A. M. Staiger, G. Ott, P. Berning, G. Lenz, D. J. Hodson, B. Kuster, R. Schmitz, H. Urlaub, M. R. Green, A. M. Melnick, R. Tooze, C. Mlynarczyk, G. Inghirami, F. Buettner, L. M. Staudt, T. Oellerich, Pathogenesis of diffuse large B cell lymphoma proteogenotypes. Cancer Cell, S1535-6108(26)00253–9 (2026).

61. J. Cox, M. Mann, MaxQuant enables high peptide identification rates, individualized p.p.b.-range mass accuracies and proteome-wide protein quantification. Nat Biotechnol 26, 1367–1372 (2008).

62. K. R. Sanson, R. E. Hanna, M. Hegde, K. F. Donovan, C. Strand, M. E. Sullender, E. W. Vaimberg, A. Goodale, D. E. Root, F. Piccioni, J. G. Doench, Optimized libraries for CRISPR-Cas9 genetic screens with multiple modalities. Nat Commun 9, 5416 (2018).

63. J. G. Doench, N. Fusi, M. Sullender, M. Hegde, E. W. Vaimberg, K. F. Donovan, I. Smith, Z. Tothova, C. Wilen, R. Orchard, H. W. Virgin, J. Listgarten, D. E. Root, Optimized sgRNA design to maximize activity and minimize off-target effects of CRISPR-Cas9. Nat Biotechnol 34, 184–191 (2016).

64. M. Wegner, K. Husnjak, M. Kaulich, Unbiased and Tailored CRISPR/Cas gRNA Libraries by SynthesizingCovalently-closed-circular (3Cs) DNA. Bio Protoc 10, e3472 (2020).

65. T. Dull, R. Zufferey, M. Kelly, R. J. Mandel, M. Nguyen, D. Trono, L. Naldini, A third-generation lentivirus vector with a conditional packaging system. J Virol 72, 8463–8471 (1998).

66. C. S. Hughes, S. Moggridge, T. Müller, P. H. Sorensen, G. B. Morin, J. Krijgsveld, Single-pot, solid-phase-enhanced sample preparation for proteomics experiments. Nat Protoc 14, 68–85 (2019).

67. V. Demichev, C. B. Messner, S. I. Vernardis, K. S. Lilley, M. Ralser, DIA-NN: neural networks and interference correction enable deep proteome coverage in high throughput. Nat Methods 17, 41–44 (2020).

68. M. E. Ritchie, B. Phipson, D. Wu, Y. Hu, C. W. Law, W. Shi, G. K. Smyth, limma powers differential expression analyses for RNA-sequencing and microarray studies. Nucleic Acids Res 43, e47 (2015).

69. P. Puigbò, I. G. Bravo, S. Garcia-Vallve, CAIcal: A combined set of tools to assess codon usage adaptation. Biol Direct 3, 38 (2008).

