## Supplementary materials for "Single-Cell Mapping of tRNA Expression Dynamics Across Human Hematopoiesis"

**The PDF file includes:**

Materials and Methods
Supplementary Tables S1 and S2
Figs. S1 to S13

**Other Supplementary Materials for this manuscript include the following:**

Supplementary Files S1 to S6

#### Materials and Methods

Bone marrow and peripheral blood sampling

Bone marrow (BM) aspirates and mobilized peripheral blood (mPB) were collected at the DRK Blood Donation Service Frankfurt from healthy volunteer donors for the purpose of unrelated allogeneic stem cell transplantation who had provided written informed consent for anonymized use of residual material for research purposes. The protocol and consent form were approved by ethics vote #329/10 of the Ethics Committee of Goethe University Medical School. Donors were healthy with no relevant past medical history, entirely normal physical examination, normal blood counts and normal biochemical laboratory workup(*40*). Donor characteristics are summarized in **Tab. S1**.

Bone marrow mononuclear cell (BMMNC) isolation

At the same day of BM aspiration, BM cells were reverse-flushed out of 150 µm in-line particle filters through which the marrow had been passed during processing for the donors with 10 mL PBS three times. BM mononuclear cells (BMMNC) were isolated from 30 mL cell suspension using density gradient centrifugation with 30 mL Pancoll human (PAN-Biotech), at 400 x g, 30 min, 30 °C, without brake. BMMNCs were frozen in freezing media containing 50 % StemSpan SFEM II (STEMCELL Technologies), 40 % FCS, 10 % DMSO, and stored below -70 °C until sc-STM-seq sample preparation.

Bulk tRNA profiling of HSPCs, progenitors and T cells

Peripheral blood mononuclear cells (PBMC) were isolated from mPB (collected by apheresis) using density gradient centrifugation as explained above. CD34+ HSPCs were MACS-enriched, and CD34+CD38low HSPCs as well as CD34+CD38high oligopotent progenitors FACS-isolated according to our established protocol(*41*). CD3+CD4+ T cells were MACS-enriched and FACS-sorted as previously described(*25*). The FACS gating strategy is depicted in **Fig. S13A**. 6,000 cells per population and donor were sorted in PBS supplemented with 500 U/mL RiboLock RNase inhibitor (Thermo Fisher), frozen, and stored below -70 °C until total RNA extraction using RNeasy Micro Kit (Qiagen) according to the manufacturer's protocol. Bulk tRNA-sequencing was executed and reads quantified according to our recently described workflow(*23*).

Principal component and unsupervised clustering analysis

Principal component analysis (PCA) and unsupervised clustering analysis of tRNA profiles of sorted HSPCs, progenitors and T cells at isotranscript level (**Fig. S13B**) as well as PCA of tRNA knockdown proteomics experiments (**Fig. S11B**) were performed using ClustVis(*42*). Unit variance scaling was applied to rows and SVD with imputation used to calculate PCs. The prediction ellipses reflect the area in which a new sample from the same group occurs with a probability of 0.95. The heatmap was generated with *average* as clustering method for rows and columns, *correlation* as clustering distance for rows, and *Manhattan* as clustering distance for columns.

sc-STM-seq library preparation and sequencing

The sc-STM-seq workflow is based on the Shasta Total RNA-seq protocol for the ICELL8 cx single-cell system (Takara). In brief, cells were fixed and long RNAs were fragmented *in situ*. Reverse transcription was then performed with the incorporation of 96 distinct barcode 1 sequences, processing ~10,000 cells per well in a 96-well plate. Following pooling, cells were redistributed into a 5,184 nano-well chip, where on-chip barcoding PCR introduced 72 x 72 barcode 2 combinations via i5 and i7 adapters.

Short and long cDNA fractions were subsequently separated, with microgel-based depletion of primer dimers. The short cDNA library (50 - 330 nt) contained reverse transcribed small RNAs, including tRNAs, whereas the long cDNA library (180 - 830 nt) predominantly comprised fragmented mRNAs and long non-coding RNAs.

The following steps were executed according to the manufacturer's protocol: thawing and fixation of BMMNCs, total RNA fragmentation and reverse transcription, on-chip cDNA amplification (PCR1), removal of rRNA-derived cDNA, and second cDNA amplification off-chip (PCR2).

The PCR2 cDNA library was then purified using SPRIselect beads (Beckman Coulter), with isolation of large cDNAs using a left-sided purification and small cDNAs using a right-sided purification. 280 uL SPRIselect beads were added to 400 uL of PCR2 product (0.7x ratio), mixed, incubated 1 min at 25 °C, vortexed at reduced speed, incubated 8 min at 25 °C and put on a magnetic rack for 5 min, followed by separation of supernatant and beads. The supernatant contained small cDNAs and was stored on ice until continuation of purification, whereas larger cDNAs were bound to the beads and immediately purified.

*Large cDNA library purification*

Beads were washed on-magnet twice with newly prepared 85% EtOH for 30 s, the pellet dried on-magnet for 1 min with open cap, 100 uL water added and the pellet rehydrated for 5 min off-magnet. The supernatant containing large cDNAs was separated after 4 min incubation on-magnet.

*Small cDNA library purification*

408 uL SPRIselect beads were added to 680 uL of supernatant containing small cDNAs (0.6x ratio), vortexed 1 min at reduced speed and incubated for 1 min at 25 °C. EtOH wash, pellet drying, rehydration and supernatant separation were executed as for the large cDNA library but the small library was eluted with 25 uL water. 20 uL of the eluate were then size-selected to contain 190 – 500 bp cDNA using the Pippin HT system (Sage Science) with a 2% agarose gel cassette.

Both libraries were pooled in a 90/10% (small/large) molar ratio and sequenced on a NovaSeq 6000 (Illumina) using the S4-300 kit, with run mode 150-8-8-150 (r1-i7-i5-r2).

<u>sc-STM-seq computational workflow</u>

*Demultiplexing and read processing*

Bcl-convert v4.4.6 (Illumina) was used to convert raw sequencing files to FASTQ files. Afterwards, the rna_demux dry run command of CogentAP and visualization in CogentDS (both Takara) were used to identify the number of expected barcodes according to the developer's protocol with the arguments --dry_run -t shasta_total_rna. CogentAP's rna_demux was then used to generate one FASTQ file per barcode according to the developer's protocol with the arguments --use_barcodes 21607 --min_reads 450 -t shasta_total_rna. Thereby, barcodes were added to each read's FASTQ header. Subsequently, all files were concatenated using the cat command in bash to one R1 and one R2 file, respectively.

Reads were trimmed using cutadapt v5.2(*43*) with the arguments -j 12 -a AGATCGGAAGAGCACACGTCTGAACTCCAGTCA -A AGATCGGAAGAGCGTCGTGTAGGGAAAGAGTGT --overlap 3 -m 1 --pair-filter=any --report=full.

Trimmed read files were uploaded to Galaxy (usegalaxy.eu)(*44*) for alignment and filtering.

*RNA workflow*

In Galaxy, reads were aligned using RNA STAR(*45*) (Galaxy Version 2.7.11b) with arguments --genomeDir Human Dec. 2013 (GRCh38/hg38) (hg38) --sjdbGTFfile Comprehensive gene annotation ALL GENCODE v49.gtf --sjdbGTFfeatureExon exon --sjdbOverhang 100 --quantMode Transcript-based BAM output (TranscriptomeSAM) --quantTranscriptomeSAMoutput prohibit indels and single-end alignments, extend softclips - compatible with RSEM --twopassMode No --chimOutType Don't report chimeric alignments --outSAMattributes NH HI AS nM NM MD ch --outSAMattrIHstart one-based --outSAMmapqUnique 60 --waspOutputMode No WASP filtering --outFilterType No --outFilterMultimapScoreRange 1 --outFilterMultimapNmax 10 --outFilterMismatchNmax 10 --outFilterMismatchNoverLmax 0.3 --outFilterMismatchNoverReadLmax 1.0 --outFilterScoreMin 0 --outFilterScoreMinOverLread 0.66 --outFilterMatchNmin 0 --outFilterMatchNminOverLread 0.66 --outSAMmultNmax -1 --outSAMtlen leftmost base of the (+)strand mate to rightmost base of the (-)mate. (+)sign for the (+)strand mate --seedSearchStartLmax 50 --seedSearchStartLmaxOverLread 1.0 --seedSearchLmax 0 --seedMultimapNmax10000 --seedPerReadNmax 1000 --seedPerWindowNmax 50 --seedNoneLociPerWindow 10 --alignIntronMin 21 --alignIntronMax 0 --alignMatesGapMax 0 --alignSJoverhangMin 5 --alignSJstitchMismatchNmax 0 --alignSJstitchMismatchNmax -1 --alignSJstitchMismatchNmax 0 --alignSJstitchMismatchNmax 0 --alignSJDBoverhangMin 3 --alignSplicedMateMapLmin 0 --alignSplicedMateMapLminOverLmate 0.66 --alignWindowsPerReadNmax 10000 --alignTranscriptsPerWindowNmax 100 --alignTranscriptsPerReadNmax 10000 --alignEndsType standard local alignment with soft-clipping allowed --peOverlapNbasesMin 0 --peOverlapMMp 0.01 --chimSegmentMin 12 --chimScoreMin 0 --chimScoreDropMax 20 --chimScoreSeparation 10 --chimScoreJunctionNonGTAG -1 --chimJunctionOverhangMin 20 --chimSegmentReadGapMax 0 --chimFilter Yes --chimMainSegmentMultNmax 10 --chimMultimapNmax 1 --chimMultimapScoreRange 1 --limitOutSJoneRead 1000 --limitOutSJcollapsed 1000000 --limitSjdbInsertNsj 1000000 --outBAMsortingBinsN 50 --winAnchorMultimapNmax 50 --outWigStrand Yes --outWigReferencesPrefix - --outWigNorm Yes.

Following STAR alignment, bam and bai files were downloaded and cell barcodes from FASTQ header added as CB tag using the script add_CB_tag.py.

To quantify read counts, featureCounts(*46*) was used with arguments -T 20 -a gencode.v49.annotation.gtf -t exon -g gene_name -s 0 -p -C -Q 0 -M -O --fraction --minOverlap 1 -R BAM.

Finally, a cell x gene matrix was created based on the XS:Z: tagged featureCounts alignment file using the Python script summarize_CB_gene_counts.py.

*tRNA workflow*

tRNAs were aligned as recently described by us(*23*). In brief, a tRNA isotranscript genome generated using the *Homo sapiens* hg38 high confidence tRNA dataset from GtRNAdb(*4*) served as a mapping reference. Adapter-trimmed R2 FASTQ files were mapped against the reference using Bowtie2(*47*, *48*) with the arguments -s 0 -u 1000000000 -5 0 -3 0 --phred33 --solexa-quals No --int-quals No -N 1 -L 10 -i S,1,0.5 --n-ceil L,0,0.15 --dpad 15 --gbar 4 --ignore-quals No --no-1mm-upfront No --local --score-min G,1,8 --ma 2 --mp 6,2 --np 0 --rdg 5 --rdg 3 --rfg 5 --rfg 3 -D 15 --seed 0 --non-deterministic No. Mapped reads were filtered to span the anticodon loop according to the GFF filtering file with BAM filter (Galaxy Version 0.5.9) and the arguments -mapped Yes -includebed [GFF filtering file]. Filtered reads were quantified using featureCounts(*46*) (Galaxy Version 2.0.3) with arguments -s Unstranded -t tRNA -g transcript_id -f No -Q 0 -M -O --fraction Yes --minOverlap 30 --fracOverlap 0 --fracOverlapFeature 0 --readExtension5 0 --readExtension3 0 -R Yes.

The generated featureCounts alignment file including XS:Z: tags was downloaded and a cell x tRNA matrix created using the Python script summarize_CB_gene_counts.py. Subsequently, tRNA isotranscript-level reads in the cell x tRNA matrix were collapsed at isodecoder level using the Python script summarize_isodecoder_to_anticodon.py to generate a cell x isodecoder matrix. Afterwards, the latter served as an input for the cell x isoacceptor matrix generation using summarize_isodecoder_to_isoacceptor.py.

*Single-cell workflow*

The cell x gene, cell x tRNA, cell x isodecoder and cell x isoacceptor matrices served as inputs for single-cell analysis in R (4.5.2) and Rstudio (2026.01.1), respectively, using the Seurat package (v. 4.3.0)(*49*).

First, a multi-layered Seurat object was created containing four assays: RNA, tRNA, isodecoder and isoacceptor. All assays were log normalized and scaled with factor 10,000. For each assay, the number of reads per cell (nCount) and the number of genes/tRNAs/isodecoders/isoacceptors per cell (nFeature) were calculated and plotted (**Fig. S2B**) as a quality control. Based on these metrics, cells were filtered to keep those with 100 – 9,000 RNA features and max. 4% mitochondrial reads. 17,788 of 21,702 cells were kept, 11,284 HD1 cells and 6,504 HD2 cells. Cells were clustered based on the RNA assay, with identification of 2,000 variable features by using vst as selection method. A principal component analysis (PCA) was computed and the standard deviation of each PC visualized as an elbow plot to identify an appropriate number of PCs to consider for Uniform Manifold Approximation and Projection (UMAP). UMAP was executed with 15 PCs and euclidean metric, resulting in 19 clusters. For each cluster, top expressed genes were identified and used for manual cell annotation based on literature research and known marker expression. In addition, cell labels were transferred from a published study(*50*) to verify cell annotations (**Fig. 1C**).

BMMNC pseudotime analysis

As differentiation trajectories of BMMNCs were not always continuous, pseudotime was investigated based on diffusion maps using the R package destiny(*51*, *52*) allowing for missing values.

A diffusion map was constructed upon extraction of PCA embeddings from the Seurat object and plotted along three diffusion components DC1, DC2, DC3 (**Fig. 2C**). Based on the diffusion map, four lineages were identified: B cell lineage, erythroid lineage, plasmacytoid dendritic cell (pDC) lineage, and monocytic lineage. Pseudotime analysis was performed with each lineage individually, meaning that the Seurat object was subset to assigned cell types and lineage-specific diffusion maps generated (**Fig. S5**). Using the 3D BMMNC diffusion map, a root and tip cell for each lineage were identified to define start and end of differentiation, and diffusion pseudotimes (DPT) were calculated and plotted onto diffusion maps.

Differentially expressed RNAs/tRNAs/isodecoders/isoacceptors along each lineage were identified using tradeSeq(*53*) (1.22.0). Downregulation of HSC- and upregulation of lineage-specific marker genes along pseudotimes verified differentiation analyses (**Fig. S6A,B**). Finally, an isodecoder synopsis figure was created as a facet plot (**Fig. S6C**).

tRNA alignment and splicing analyses

In Galaxy, the filtered tRNA BAM file was converted using BAM-to-SAM(*54*) to allow extraction of the alignment length by summarizing the arguments M, D, N and =. Summed read counts were plotted against the alignment length using ggplot2 (**Fig. 1B**).

To perform pseudobulk alignment for each cluster, corresponding barcodes were csv exported from the Seurat object using R. The cluster-barcode lists enabled subsequent export of alignments per cluster in bash. The generated cluster-BAM files enabled inspection of read coverage and tRNA splicing using the Integrative Genomics Viewer (IGV) for Windows, version 2.19.1(*55*) (**Fig. S4C**).

For splicing quantification, the average read count of the three exonic nucleotides upstream (ex1) and downstream (ex2) of the intron were divided by the average read count of intronic nucleotides. This spliced/unspliced ratio was calculated for each cluster (cell type) with at least one read at each intronic nucleotide, and plotted using GraphPad Prism 11.0.1 (**Fig. S4B**).

Re-clustering of the HSPC compartment

The generated Seurat object was subset to HSPCs, normalized, scaled, variable features identified and a PCA computed as explained above. UMAP was executed on the RNA assay, top expressed genes identified and cell types annotated considering our recently published single-cell HSPC dataset(*25*) (**Fig. 3B**).

Differential tRNA expression analyses

DEseq2 (1.40.2)(*56*) was used for differential tRNA expression analysis of sc-STM-seq pseudobulk comparisons and sorted CD34+CD38low HSPCs compared to CD3+CD4+ T cells. Read counts were extracted per cluster (cell type) and donor to create volcano plots using ggplot2 in R (**Fig. 2A,B, 3A,C**, **S4A**, and **S10C**). DEseq2 results are summarized in **supplementary files S1, S2,** and **S4**.

Comparison of sc-STM-seq with bulk tRNA-seq

To compare sc-STM-seq with an established bulk tRNA-seq workflow, raw tRNA counts were extracted from the sc-STM-seq Seurat object in R and plotted against bulk tRNA-seq raw counts from our previously published dataset comprising whole PB from 10 healthy donors(*23*) (**Fig. S2A**). Spearman correlations were calculated using GraphPad Prism 11.0.1.

In addition, log normalized sc-STM-seq tRNA counts of HSPCs (pseudobulk of cluster 7) were compared with bulk tRNA-seq tpm normalized counts of sorted mPB-derived CD34+ HSPCs. The same comparison was executed with CD4+ T cells (pseudobulk of cluster 0) and sorted mPB-derived CD3+CD4+ T cells. Normalized tRNA expressions from sc-STM-seq were extracted from the Seurat object using R, Spearman correlations and plots were created with GraphPad Prism 11.0.1 (**Fig. S2A**).

Comparison of sc-STM-seq tRNA and mRNA expression

A violin plot was created in R comparing the log-transformed expression level and number of expressing cells of three tRNAs and three mRNAs. A highly expressed, intermediately expressed and lowly expressed tRNA and mRNA, respectively, were chosen (**Fig. S3**).

tRNA expression dot plots

Dot plots indicating isodecoder/isoacceptor expression levels and percentage of expressing cells were created in R using ggplot2. (**Fig. S7** and **S8**).

tRNA-based clustering analysis

To cluster cells based on their tRNA expression profiles, Seurat was used and the default assay was set to "tRNA" instead of "RNA". Data normalization, scaling and PCA were thereby based on tRNA expression. UMAP was conducted including 10 PCs resulting in 14 clusters (**Fig. 4A**). Using R, a stacked bar plot was generated visualizing the cell type proportions (annotated based on RNA expression) for each tRNA cluster (**Fig. 4B**). The same workflow was used to perform isodecoder- and isoacceptor-based clustering, including UMAP and stacked bar plot generation. For better readability, cell proportions per tRNA-cluster were exported and visualized as bar plots for selected cell types using GraphPad Prism 11.0.1 (**Fig. 4C**).

Data preparation for web-based atlas display

The web-based tRNA expression atlas can be accessed via https://bioinformatics.mpi-bn.mpg.de/hematopoietic-tRNA-atlas.

The previously described matrices for tRNA and RNA expression were used as input for the SC framework(*57*) and transferred into an AnnData object(*58*). The data were subsequently processed and filtered to ensure compatibility with the Cellxgene VIP application(*59*).

For low-dimensional representation of cellular heterogeneity, both principal component analysis (PCA) and uniform manifold approximation and projection (UMAP) embeddings were incorporated into the final object. To enable integrated exploration of transcriptomic and tRNA information, RNA and tRNA count matrices were combined into a single joined matrix representation. Feature identities were preserved through modality-specific prefixes, allowing RNA and tRNA expression values to be queried and visualized independently within the application environment.

The application is hosted within the CPI Translational Hub2 infrastructure, leveraging a cloud-native deployment architecture based on Kubernetes orchestration and Docker container virtualization to ensure scalability, reproducibility, and permanent public availability.

Computational resources are provided by the Max Planck Institute for Heart and Lung Research and the German Network for Bioinformatics Infrastructure (deNBi).

Quantitative proteomics of primary HSPCs and T cells

$1x10^6$ CD34+ HSPCs and $1x10^6$ CD3+CD4+ T cells from each of four healthy stem cell donors were MACS-enriched from mPB, pelleted, snap-frozen and stored below -70 °C until sample preparation. Quantitative proteomics experiments were basically conducted as described in detail previously(*60*). Briefly, cell pellets were lysed in 50 µl Urea buffer (8 M urea, 20 mM HEPES, pH 8.0, 1 mM sodium orthovanadate, 2.5 mM sodium pyrophosphate, 1 mM beta-glycerophosphate), followed by reduction/alkylation of extracted proteins and digestion with Lys-C (Wako) and Trypsin (Promega). Protein lysate preparation failed for the CD34+ sample of donor 1. For relative protein quantification, 10 µg of each peptide sample were individually labeled with 100 µg of tandem mass tags (TMT) according to the instructions of the manufacturer (Thermo Fisher Scientific) and combined in a multiplexed sample. The peptide mixture was pre-fractionated using High pH RPLC kit (Thermo Fisher Scientific) and the eight individual fractions were analyzed by LC-MS/MS on an Ultimate 3000 HPLC system coupled to a Q Exactive HF quadrupole-Orbitrap hybrid mass spectrometer via a Nanospray Flex electrospray ionization source (all Thermo Fisher Scientific). MS raw data was processed with the MaxQuant software(*61*) and for protein identification the mass spectra were searched against the Uniprot human reference proteome using the integrated Andromeda engine. For protein quantitation, the TMT reporter ion intensities were extracted on the MS2 level and adjusted for equal peptide loading. Results of the analysis are summarized in **supplementary file S3**. Volcano plots with padj cut-off of 0.05 and annotation of selenoproteins (**Fig. S9**) or LMNB2, TALDO1, and PRDX1 (**Fig. S10B**) were created using R.

tRNA knockdown experiments

*Cell culture*

The pro-B cell line SEM (ACC 546**)** lentivirally transduced to express *Streptococcus pyogenes* Cas9 and a blasticidin resistance was kindly provided by Anjali Cremer, Goethe University Hospital Frankfurt. SEM cells were maintained in IMDM supplemented with 10 % FBS, 2 mM L-glutamine, 100 U/ml penicillin, 100 µg/ml streptomycin, and 10 ug/mL blasticidin, under sterile conditions. Cells were cultivated at 37 °C in 5% $CO_2$ and a relative humidity of 95%. Passaging was performed twice a week to maintain a cell density of $1\text{-}3x10^6$ cells/mL. Lenti-X HEK293T cells (Takara Bio, 632180) were maintained as adherent culture in DMEM supplemented with 10 % FBS, 2 mM L-glutamine, 100 U/ml penicillin, and 100 µg/ml streptomycin, under sterile conditions. Cells were cultivated at 37 °C in 5% $CO_2$ and a relative humidity of 95%. Cells were passaged three times per week with complete media exchange through detachment using trypsin after washing with PBS.

*gRNA design and cloning*

gRNAs were designed using the CRISPick design tool(*62*, *63*) for targeting tRNA-Asp-GTC-1 (gRNA-1: ACCGGGGTTCAATTCCCCGA; gRNA-2: TAGTATCCCCGCCTGTCACG), tRNA-Asp-GTC-2 (gRNA-1: GAGTATCCCCGCCTGTCACG; gRNA-2: ACCGGGGTTCGATTCCCCGA), tRNA-Glu-TTC-2 (gRNA-1: CCCGGGTTCGACTCCCGGTG; gRNA-2: GAGTCGAACCCGGGCCGCCT), and tRNA-Glu-CTC-1 and -2 (gRNA-1: CGCTCTCACCGCCGCGGCCC; gRNA-2: GTGGTCTAGTGGTTAGGATT). gRNAs were inserted into the 3Cs expression vector (Addgene: 189632) by the 3Cs cloning strategy as explained before(*64*). gRNA-containing 3Cs oligonucleotides were obtained from Integrated DNA Technologies (IDT). The non-targeting control gRNA (AATTAATTAATGTTAATTAA) was used as baseline control. Each plasmid was validated by Sanger sequencing (Microsynth AG).

*Virus production*

The cloned plasmids were encapsulated in vesicular stomatitis virus G (VSVG)-pseudotyped lentiviral particles using calcium phosphate transfection of Lenti-X HEK293T cells and a split-genome strategy(*65*). Concentration of virus particles was executed by ultracentrifugation at 50,000 x g for 1 h at 4 °C. Subsequently, titers were quantified via limited dilution transduction of SEM-Cas9 cells and viability assessment following 72 h of puromycin treatment (2 ug/mL). gRNA expression constructs were stably integrated into SEM-Cas9 cells by lentiviral transduction with a multiplicity of infection of 0.15.

*tRNA-seq and knockdown efficiency calculation*

Transduced cells were selected under puromycin treatment (2 ug/mL) for 72 h, followed by pelleting of $1x10^6$ cells and total RNA extraction using the RNeasy Mini Kit (Qiagen, 74104) according to the manufacturer's protocol. Bulk tRNA-sequencing was executed and reads quantified according to our recently described workflow(*23*) and knockdown efficiencies calculated as 1 minus the ratio of knockdown to NTC mean tpm values. DEseq2 (1.40.2)(*56*) was used for differential tRNA expression analysis and a volcano plot created using ggplot2 in R (**Fig. S11A**). Raw data, knockdown efficiencies, and DEseq2 results are summarized in **supplementary file S5**.

*Quantitative proteomics of tRNA knockdown samples*

Transduced cells were selected under puromycin treatment (2 ug/mL) for 72 h, followed by pelleting and snap-freezing of $1.2x10^6$ cells. Cell pellets were stored below -70 °C until protein lysate generation.

*Proteomics Sample Preparation*

Frozen pellets were resuspended in 500 µL lysis buffer (0.5% n-Dodecyl β-D-maltopyranoside in 20 mM HEPES, pH 8.0, 1 mM sodium orthovanadate, 2.5 mM sodium pyrophosphate, 1 mM beta-glycerophosphate), the suspension was sonicated two times for 10 s using a probe sonicator on ice and protein concentrations of the cleared lysates were determined using Pierce A660 protein assay (Thermo Fisher Scientific). Samples with equal volumes according to 50 µg of protein were supplemented with 500 U/mL benzonase (Merck) and incubated for 30 min on ice. Subsequently, samples were reduced with Tris(2-carboxyethyl)phosphine (TCEP, 10 mM for 30 min at 37 °C) and alkylated with iodoacetamide (IAA, 25 mM for 15 min at 37 °C in the dark). Proteins were cleaned up using an adapted version of the SP3 protocol(*66*): hydrophilic and hydrophobic carboxylated, paramagnetic beads (Cytiva) were mixed 1:1, washed three times with water to remove the storage buffer and added in a protein:bead (w/w) ratio of 1:10 to each sample. Protein binding was induced by the addition of pure acetonitrile (ACN) to a final concentration of 70% and the sample-bead-mixture was incubated at RT for 18 min, shaking at 850 rpm. Samples were placed on a magnet and left to settle for 2 min after which the supernatant was discarded. Samples were then washed two times with 80% ethanol and once with pure ACN, washing supernatant being discarded each time. Proteins were eluted from the beads by adding 20 mM HEPES (pH 8.0) and were digested overnight with trypsin (MS grade, Promega) at 37 °C and 1:50 (w/w) enzyme-to-substrate ratio. The peptide mixtures were acidified with trifluoroacetic acid (TFA), purified on an Agilent Bravo liquid handler using C18 cartridges (AssayMAP 5 µl, Agilent) and dried by vacuum centrifugation.

*LC-MS Measurement*

The dried peptide samples were dissolved in 0.1% formic acid (FA) and peptide concentrations were determined using Pierce fluorometric assay (Thermo Fisher Scientific). The peptide samples were analyzed by LC-MS/MS on a Vanquish Neo UHPLC system coupled to an Orbitrap Astral Zoom mass spectrometer via a Nanospray Flex electrospray ion source and a FAIMS interface (all Thermo Fisher Scientific) using a data-independent acquisition scheme

(DIA). Per run, 200 ng of peptides were concentrated and desalted on a NanoShield C18 trap column (pore size 120 Å, particle size 3 µm, inner diameter 100 µm, length 50 mm, IonOpticks), followed by separation at 50 °C on a 25 cm Aurora Ultimate C18 analytical column (pore size 120 Å, particle size 1.7 µm, inner diameter 75 µm, IonOpticks) using a 25 min method with a 20 min active gradient of 2% to 40% solvent B (0.1% FA in ACN) over solvent A (0.1% FA) at a flow rate of 400 nL/min. Precursor ion survey scans were acquired using the Orbitrap mass analyzer with the following parameters: resolution 240,000, scan range *m/z* 380-980, automatic gain control (AGC) target $5x10^6$, maximum injection time 3 ms, RF lens setting 40% and FAIMS compensation voltage -45 V. For fragment ion scans using the Astral mass analyzer in the range of *m/z* 150-2000, precursor ions were isolated for collision-induced dissociation (HCD) with an isolation window of 2 Th width, resulting in 300 MS/MS scan events through each survey scan. The normalized HCD collision energy was set to 25% and for fragment ion analysis the AGC target value was $1x10^4$ at a maximum injection time of 3 ms.

*Data Base Search*

Raw DIA data were analyzed using the DIA-NN software (version 2.6.1 Academia)(*67*) and spectra were searched against the Uniprot human proteome (release 2025_04) and 245 frequently observed contaminants, using a spectral library predicted by DIA-NN. The mass tolerance for precursor ions was set to 4 ppm and for fragment ions to 10 ppm, oxidation of methionine was considered as dynamic modification and carbamidomethylation of cysteine was defined as fixed modification. The peptide length was considered 7 to 30 amino acid residues with up to three missed cleavage sites allowed. Match between runs was activated and the maximum false discovery rate (FDR) was set to 5 %.

*Statistical Analysis*

The output data was filtered by removing potential contaminants. To control for equal sample loading, relative protein abundances from each LC-MS/MS run were normalized on the median of the summed-up intensities from each sample. The dataset was filtered for a maximum missing rate of 30 % and differential expression analysis between replicate groups was conducted using limma(*68*) with a maximum FDR of 5 %. Raw data and differential expression analysis results are summarized in **supplementary file 6**.

*Visualization*

Raw data were analyzed using a PCA created with ClustVis(*42*). As before, unit variance scaling was applied to rows and SVD with imputation used to calculate PCs. The prediction ellipses indicate the area in which a new sample from the same group occurs with probability 0.95. Differential expression results were visualized using volcano plots with R (**Fig. S11C, D**) and a scatter plot (**Fig. S11E**) generated by plotting the log2FC obtained under DMSO treatment against the corresponding log2FC obtained under MG132 treatment, with each point representing one protein. Protein significance was determined separately for each condition using adjusted P values from the differential expression analysis. Proteins were classified as significantly altered under DMSO only, MG132 only, both conditions, or neither condition according to an adjusted P value threshold of 0.05. To visualize statistical confidence, the significance score displayed in the plot was calculated as the minimum of the two negative log10-transformed adjusted P values, reflecting the weaker statistical support across the two conditions.

*Amino acid composition analysis*

Protein amino acid sequences were retrieved from UniProt (release 2025_04) using the corresponding UniProt accession of each quantified protein. For each protein, the relative abundance of each of the 20 canonical amino acids was calculated as the fraction of residues corresponding to a given amino acid divided by the total protein length. Proteins were classified as upregulated, downregulated, or not differentially expressed based on the results of the proteomic differential expression analysis for the comparison DMSO-AspGluKD vs. DMSO-NTC. For each amino acid, distributions of amino acid fractions among the three groups were compared using a Kruskal-Wallis test. P values were adjusted for multiple testing across all amino acids using the Banjamini-Hochberg procedure. Pairwise comparisons between groups were performed using two-sided Wilcoxon rank-sum tests only for amino acids with a significantly adjusted Kruskal-Wallis test (adjusted $P < 0.05$). Pairwise P values were corrected using the Holm method. Individual proteins were visualized as jittered points overlaid on boxplots displaying the median and interquartile range; for visualization purposes, a random subset of proteins from the non-differentially expressed groups was plotted when necessary to reduce overplotting, whereas all differentially expressed proteins were displayed (**Fig. S12C, D**). Results of these analyses are summarized in **supplementary file 6**.

<u>Codon usage analyses</u>

The coding sequences of the canonical transcripts of the investigated genes (*LMNB2* (ENST00000325327.4), *TALDO1* (ENST00000319006.8), *PRDX1* (ENST00000319248.13, or those identified in the differential protein expression analysis upon tRNA knockdown) were manually retrieved from ensembl.org and their codon usages calculated as codons per million (cpm) using the CAIcal online tool(*69*). The human genomic codon usage was retrieved from CAIcal as well, and GraphPad Prism version 11.0.1 was used to visualize both parameters in an XY coordinate system, calculate Spearman correlation factors, and simple linear regressions (**Fig. S10A** and **S12A, B**).

<u>Statistical analyses</u>

If not otherwise specified, statistical analyses were performed using R (DEseq2) or GraphPad Prism (Spearman correlation testing) version 11.0.1 for Windows, GraphPad Software, Boston, Massachusetts USA, www.graphpad.com. DEseq2 results are summarized in **supplementary files S1**, **S2**, **S4**, and **S5**.

**Table S1.**
Donor characteristics.

| **Experiment** | **Donor** | **Source** | **Age at donation** | **Sex** |
|---|---|---|---|---|
| sc-STM-seq | HD1 | Bone marrow | 28 | Male |
| | HD2 | Bone marrow | 27 | Female |
| bulk tRNA-seq | D1 | Mobilized peripheral blood | 30 | Male |
| | D2 | Mobilized peripheral blood | 32 | Male |
| | D3 | Mobilized peripheral blood | 24 | Male |
| quantitative proteomics | D1 | Mobilized peripheral blood | 30 | Male |
| | D2 | Mobilized peripheral blood | 51 | Female |
| | D3 | Mobilized peripheral blood | 29 | Male |
| | D4 | Mobilized peripheral blood | 41 | Male |

5

**Table S2.**

tRNA and RNA read counts per cell type. Note that a tRNA read resembles a tRNA molecule, whereas RNA reads resemble only tiny fragments of an mRNA or long non-coding RNA.

| Cluster | Cell type | Cell number | Median tRNA reads/cell | Range tRNAs/cell | Median RNA reads/cell | Range RNA reads/cell |
|---|---|---|---|---|---|---|
| 0 | CD4+ T cells | 2,563 | 358 | 1 - 7,381 | 123,270 | 4,099 - 557,416 |
| 1 | Monocytes | 2,559 | 388 | 1 - 6,051 | 164,837 | 21,159 - 1,158,053 |
| 2 | CD8+ T cells | 2,200 | 506 | 1 - 5,584 | 133,217 | 3,373 - 533,576 |
| 3 | Myelocytes | 1,616 | 311 | 1 - 5,134 | 90,246.5 | 4,651 - 651,124 |
| 4 | pDC | 1,274 | 1,514 | 1 - 6,320 | 234,375 | 28,665 - 731,097 |
| 5 | GMP | 1,049 | 540 | 1 - 12,068 | 287,761 | 28,510 - 1,807,720 |
| 6 | Mature B cells | 946 | 392 | 1 - 5,787 | 130,654.5 | 23,066 - 903,732 |
| 7 | HSPCs | 903 | 861 | 3 - 14,457 | 366,384 | 34,064 - 1,543,154 |
| 8 | cDC 2 | 892 | 1,225.5 | 1 - 8,652 | 268,900 | 64,438 - 1,096,342 |
| 9 | NK cells | 772 | 528.5 | 1 - 17,392 | 144,716 | 27,780 - 1,077,680 |
| 10 | Late erythroid progenitors | 731 | 262 | 1 - 6,606 | 126,013 | 10,502 - 783,526 |
| 11 | Granulocytes | 611 | 53 | 1 - 2,071 | 101,626 | 23,631 - 419,881 |
| 12 | Erythroid cells | 537 | 236 | 1 - 3,348 | 81,152 | 22,970 - 550,329 |
| 13 | Pre-B/immature B cells | 393 | 395 | 1 - 5,050 | 155,190 | 22,240 - 1,474,716 |
| 14 | Early erythroid progenitors | 338 | 679.5 | 2 - 12,576 | 493,340.5 | 70,514 - 1,714,003 |
| 15 | Plasma cells | 168 | 359 | 3 - 8,839 | 273,094.5 | 62,347 - 693,706 |
| 16 | Pro-B cells | 123 | 311 | 2 - 7,330 | 201,498 | 29,381 - 792,256 |
| 17 | cDC 1 | 81 | 1,948 | 7 - 5,567 | 338,833 | 102,155 - 616,164 |
| 18 | Mesench./strom. cells | 32 | 689 | 7- 8,779 | 343,565 | 31,265 - 1,433,721 |

**Supplementary figures**

**Figure S1**

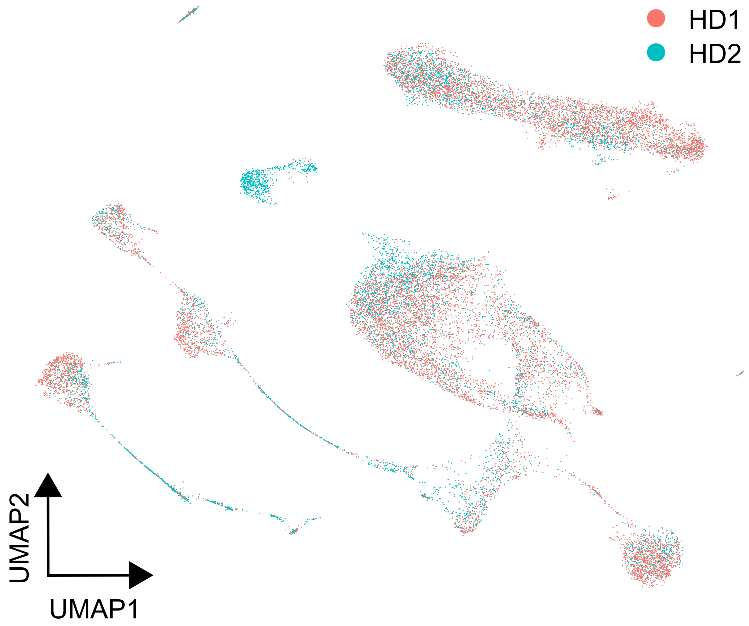

**Fig. S1. Distribution of cells by donors.**
A UMAP indicates which cells were derived from which donor. HD healthy donor.

**Figure S2**

A

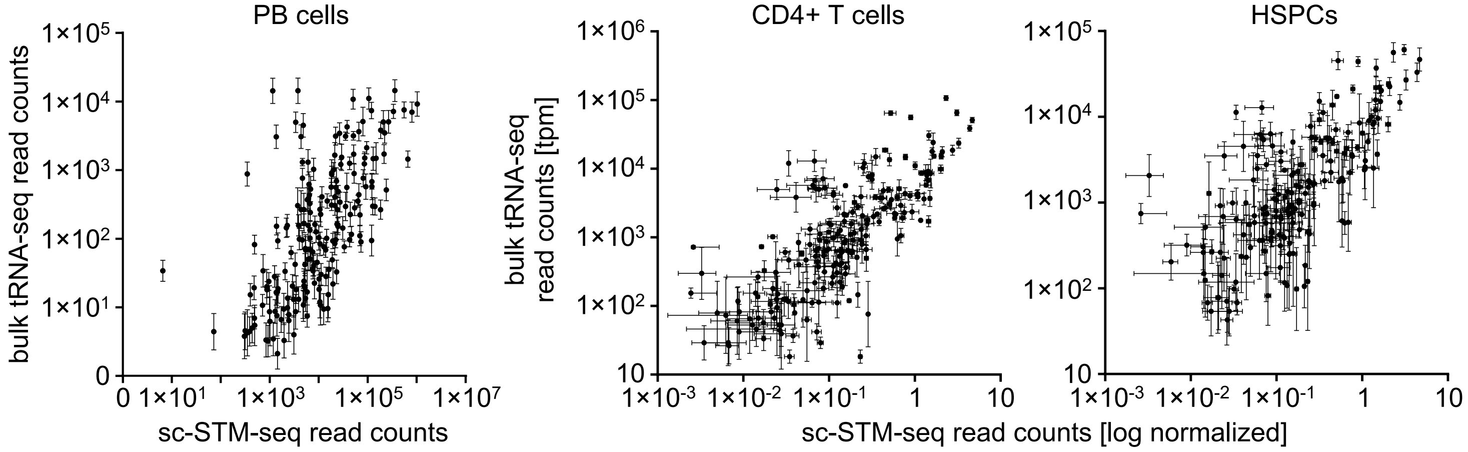

| | PB cells | CD4+ T cells | HSPCs |
|---|---|---|---|
| Spearman r | 0.8117 | 0.8425 | 0.8220 |
| 95% confidence interval | 0.7646 to 0.8501 | 0.8023 to 0.8751 | 0.7772 to 0.8585 |
| P value | | | |
| P (two-tailed) | <0.0001 | <0.0001 | <0.0001 |
| P value summary | **** | **** | **** |
| Exact or approx. P value? | Approximate | Approximate | Approximate |
| Significant? (alpha = 0.05) | Yes | Yes | Yes |
| Number of XY Pairs | 265 | 265 | 265 |

B

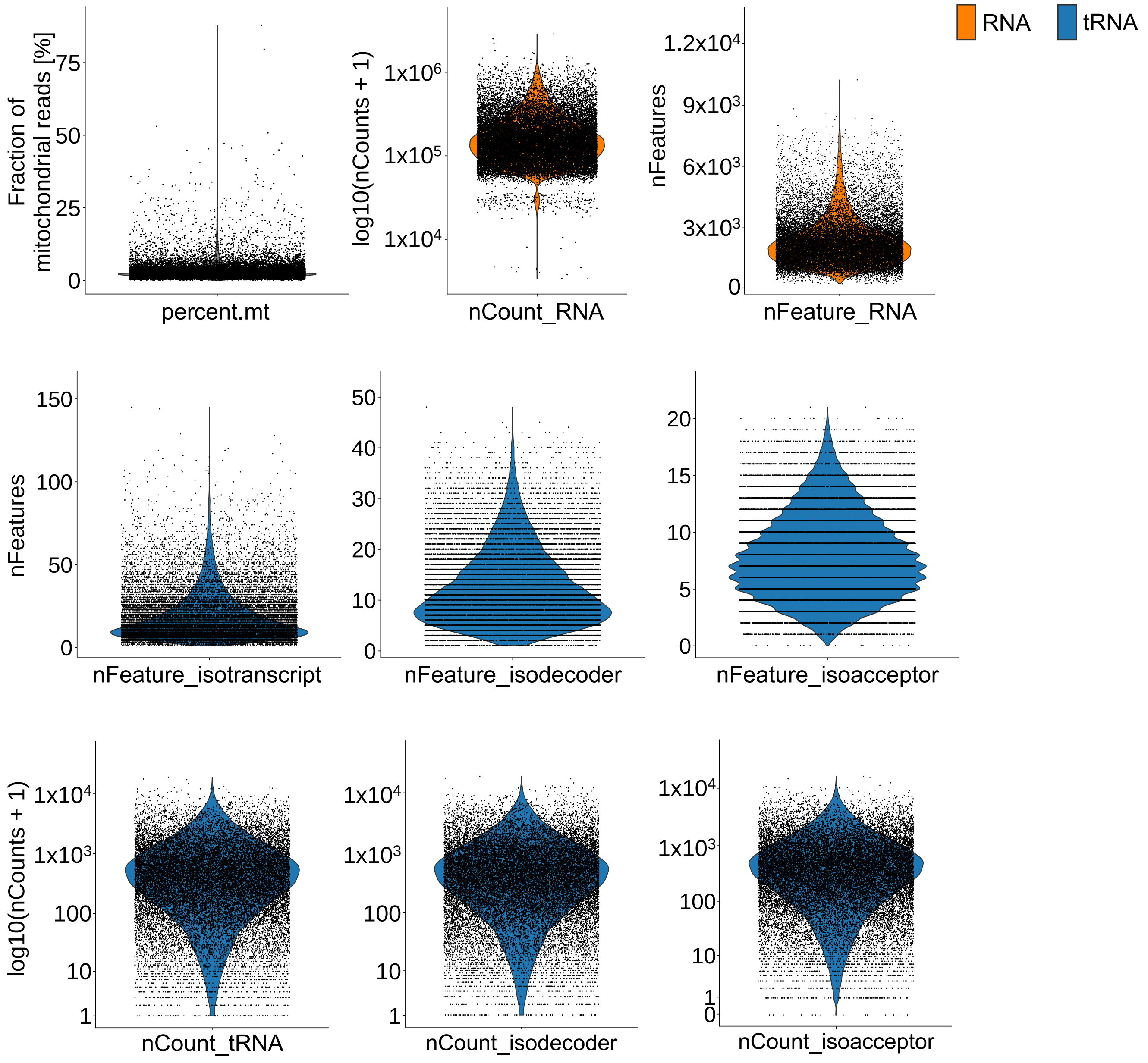

**Fig. S2. sc-STM-seq quality control parameters.**

**A** Spearman correlation testing of sc-STM-seq read counts against bulk tRNA-seq counts of PB cells(*23*), as well as prospectively FACS-isolated CD4+ T cells and CD34+ HSPCs, respectively. **B** Violin plots indicating all quality control parameters assessed in Seurat of the RNA (orange) and tRNA (blue) assays before filtering of cells.

**Figure S3**

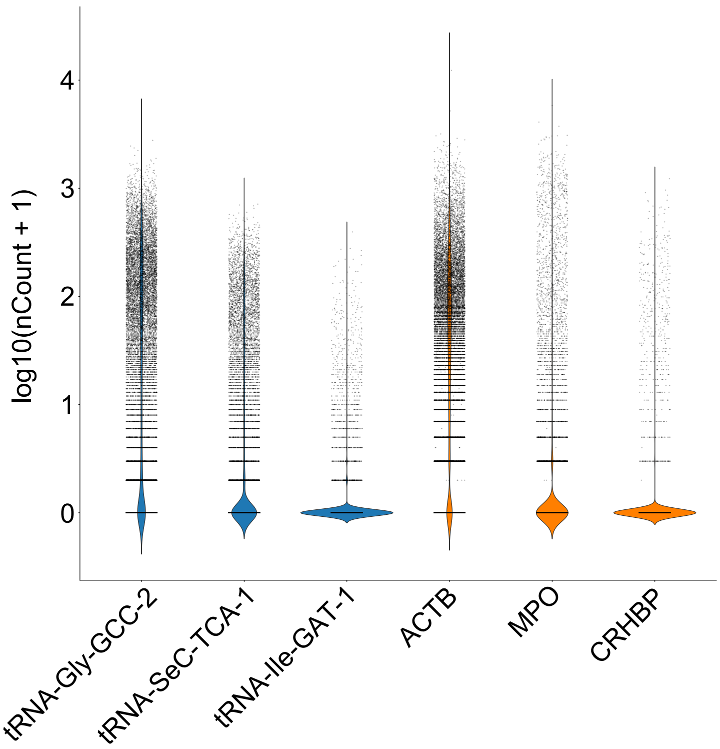

log10(nCount + 1)
4
3
2
1
0
tRNA-Gly-GCC-2
tRNA-SeC-TCA-1
tRNA-Ile-GAT-1
ACTB
MPO
CRHBP

**Fig. S3. sc-STM-seq tRNA and mRNA expression comparison.**
Violin plots comparing tRNA and mRNA expression distribution of a highly (tRNA-Gly-GCC-2, ACTB), intermediately (tRNA-SeC-TCA-1, MPO), and lowly (tRNA-Ile-GTA-1, CRHBP) expressed tRNA and mRNA, respectively.

### Figure S4

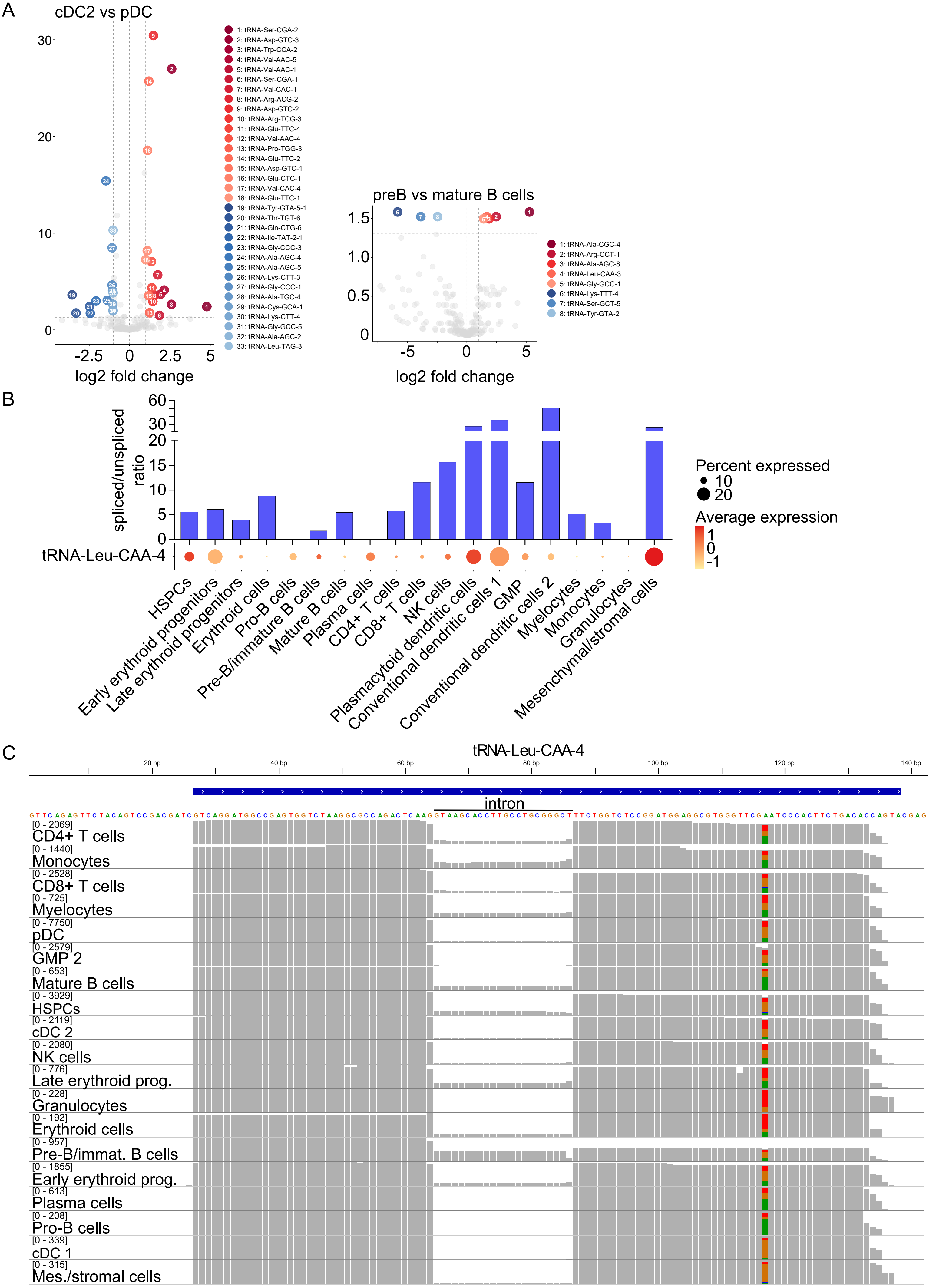

A
cDC2 vs pDC
log2 fold change
1: tRNA-Ser-CGA-2
2: tRNA-Asp-GTC-3
3: tRNA-Trp-CCA-2
4: tRNA-Val-AAC-5
5: tRNA-Val-AAC-1
6: tRNA-Ser-CGA-1
7: tRNA-Val-CAC-1
8: tRNA-Arg-ACG-2
9: tRNA-Asp-GTC-2
10: tRNA-Arg-TCG-3
11: tRNA-Glu-TTC-4
12: tRNA-Val-AAC-4
13: tRNA-Pro-TGG-3
14: tRNA-Glu-TTC-2
15: tRNA-Asp-GTC-1
16: tRNA-Glu-CTC-1
17: tRNA-Val-CAC-4
18: tRNA-Glu-TTC-1
19: tRNA-Tyr-GTA-5-1
20: tRNA-Thr-TGT-6
21: tRNA-Gln-CTG-6
22: tRNA-Ile-TAT-2-1
23: tRNA-Gly-CCC-3
24: tRNA-Ala-AGC-4
25: tRNA-Ala-AGC-5
26: tRNA-Lys-CTT-3
27: tRNA-Gly-CCC-1
28: tRNA-Ala-TGC-4
29: tRNA-Cys-GCA-1
30: tRNA-Lys-CTT-4
31: tRNA-Gly-GCC-5
32: tRNA-Ala-AGC-2
33: tRNA-Leu-TAG-3
preB vs mature B cells
log2 fold change
1: tRNA-Ala-CGC-4
2: tRNA-Arg-CCT-1
3: tRNA-Ala-AGC-8
4: tRNA-Leu-CAA-3
5: tRNA-Gly-GCC-1
6: tRNA-Lys-TTT-4
7: tRNA-Ser-GCT-5
8: tRNA-Tyr-GTA-2
B
spliced/unspliced ratio
tRNA-Leu-CAA-4
Percent expressed
10
20
Average expression
HSPCs
Early erythroid progenitors
Late erythroid progenitors
Erythroid cells
Pro-B cells
Pre-B/immature B cells
Mature B cells
Plasma cells
CD4+ T cells
CD8+ T cells
NK cells
Plasmacytoid dendritic cells
Conventional dendritic cells 1
GMP
Conventional dendritic cells 2
Myelocytes
Monocytes
Granulocytes
Mesenchymal/stromal cells
C
tRNA-Leu-CAA-4
intron
[0 - 2069] CD4+ T cells
[0 - 1440] Monocytes
[0 - 2528] CD8+ T cells
[0 - 725] Myelocytes
[0 - 7750] pDC
[0 - 2579] GMP 2
[0 - 653] Mature B cells
[0 - 3929] HSPCs
[0 - 2119] cDC 2
[0 - 2080] NK cells
[0 - 776] Late erythroid prog.
[0 - 228] Granulocytes
[0 - 192] Erythroid cells
[0 - 957] Pre-B/immat. B cells
[0 - 1855] Early erythroid prog.
[0 - 613] Plasma cells
[0 - 208] Pro-B cells
[0 - 339] cDC 1
[0 - 315] Mes./stromal cells

**Fig. S4. sc-STM-seq data exploration.**

**A** Extended differential tRNA expression analysis between indicated cell types. **B** Splicing efficiency of tRNA-Leu-CAA-4 quantified as the ratio of spliced to unspliced reads for all cell types. The dot plot indicates average tRNA expression and proportion of cells expressing the tRNA. **C** tRNA alignment bam coverage of all cell types in pseudobulk. The tRNA sequence incl. the intron is indicated in blue, with pseudoadapters up- and downstream. The relative alignment quantification is indicated as grey bars, with absolute read counts indicated on the left-hand side. For many cell types, unprocessed tRNA reads including the intron were detected. A misincorporation at A58 was detected in all cell types.

**Figure S5**

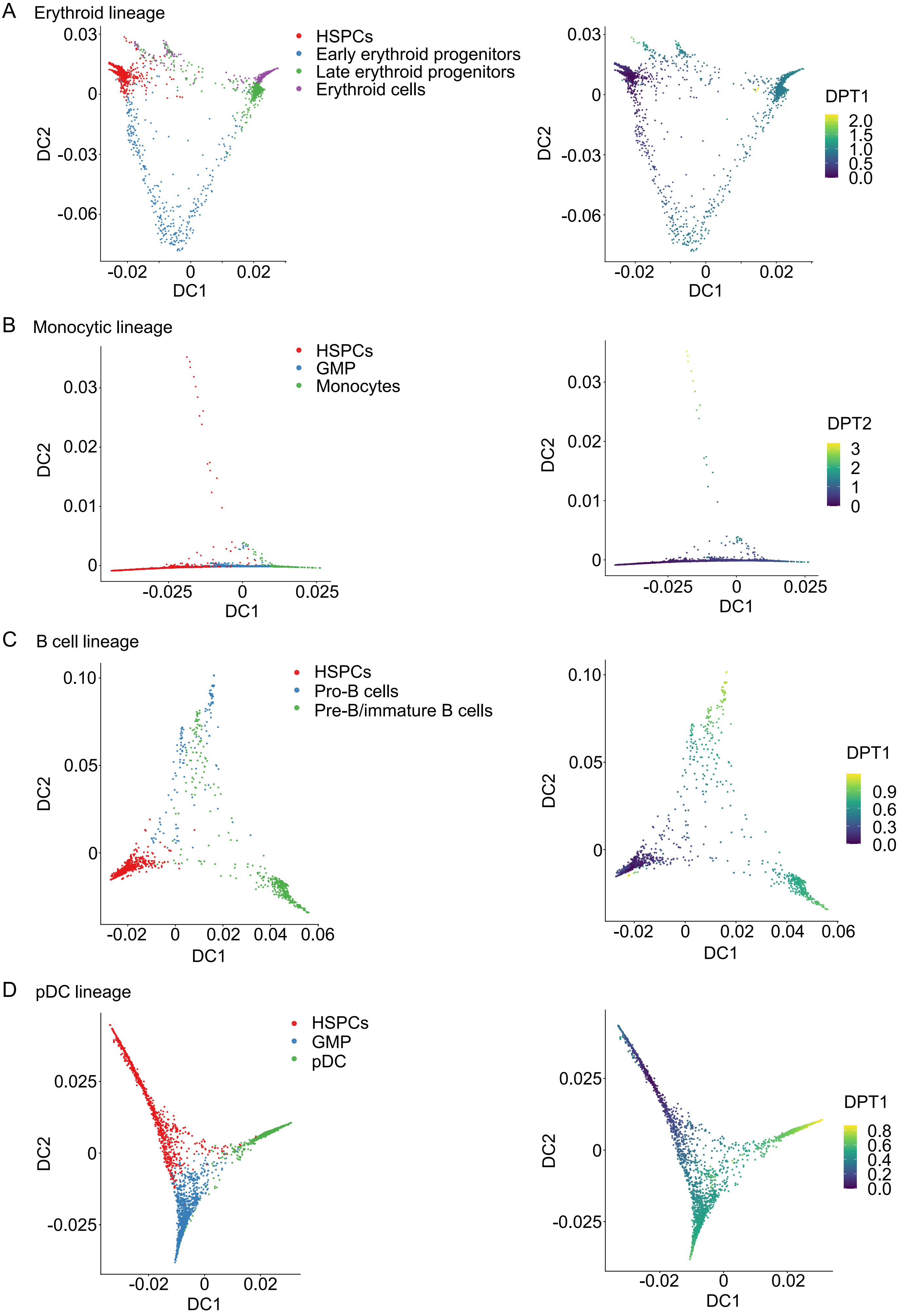
A
Erythroid lineage
HSPCs
Early erythroid progenitors
Late erythroid progenitors
Erythroid cells
DPT1
DC1
DC2
B
Monocytic lineage
HSPCs
GMP
Monocytes
DPT2
DC1
DC2
C
B cell lineage
HSPCs
Pro-B cells
Pre-B/immature B cells
DPT1
DC1
DC2
D
pDC lineage
HSPCs
GMP
pDC
DPT1
DC1
DC2

**Fig. S5. Diffusion maps of individual lineages.**

Diffusion maps were generated including the indicated cell types and diffusion pseudotimes (DPT) plotted onto diffusion maps. **A** Erythroid lineage. **B** Monocytic lineage. **C** B cell lineage. **D** Plasmacytoid dendritic cell (pDC) lineage.

### Figure S6

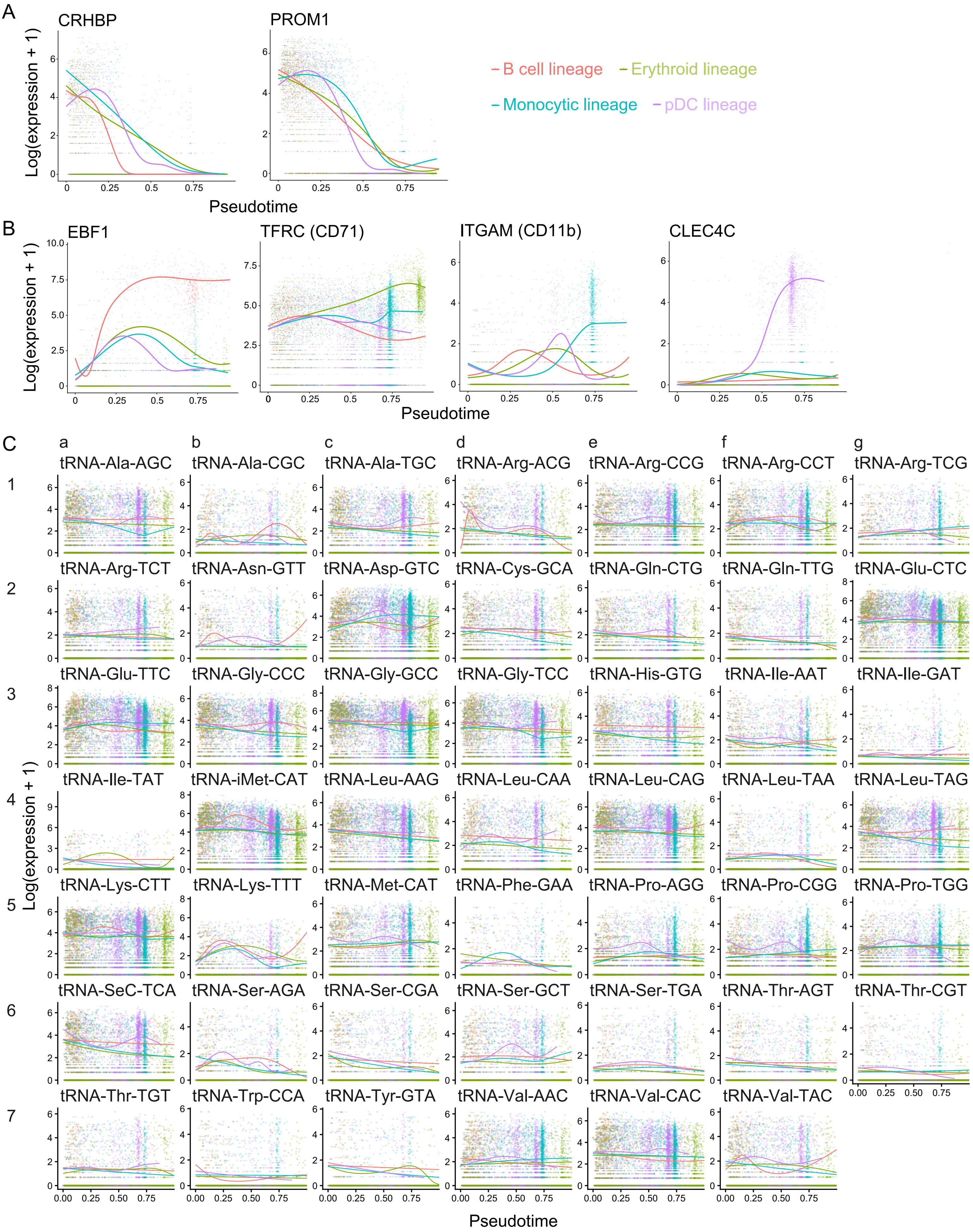
A
CRHBP
PROM1
Log(expression + 1)
Pseudotime
B cell lineage
Erythroid lineage
Monocytic lineage
pDC lineage
B
EBF1
TFRC (CD71)
ITGAM (CD11b)
CLEC4C
Log(expression + 1)
Pseudotime
C
a
b
c
d
e
f
g
1
2
3
4
5
6
7
tRNA-Ala-AGC
tRNA-Ala-CGC
tRNA-Ala-TGC
tRNA-Arg-ACG
tRNA-Arg-CCG
tRNA-Arg-CCT
tRNA-Arg-TCG
tRNA-Arg-TCT
tRNA-Asn-GTT
tRNA-Asp-GTC
tRNA-Cys-GCA
tRNA-Gln-CTG
tRNA-Gln-TTG
tRNA-Glu-CTC
tRNA-Glu-TTC
tRNA-Gly-CCC
tRNA-Gly-GCC
tRNA-Gly-TCC
tRNA-His-GTG
tRNA-Ile-AAT
tRNA-Ile-GAT
tRNA-Ile-TAT
tRNA-iMet-CAT
tRNA-Leu-AAG
tRNA-Leu-CAA
tRNA-Leu-CAG
tRNA-Leu-TAA
tRNA-Leu-TAG
tRNA-Lys-CTT
tRNA-Lys-TTT
tRNA-Met-CAT
tRNA-Phe-GAA
tRNA-Pro-AGG
tRNA-Pro-CGG
tRNA-Pro-TGG
tRNA-SeC-TCA
tRNA-Ser-AGA
tRNA-Ser-CGA
tRNA-Ser-GCT
tRNA-Ser-TGA
tRNA-Thr-AGT
tRNA-Thr-CGT
tRNA-Thr-TGT
tRNA-Trp-CCA
tRNA-Tyr-GTA
tRNA-Val-AAC
tRNA-Val-CAC
tRNA-Val-TAC
Log(expression + 1)
Pseudotime

**Fig. S6. Extended Tradeseq analysis.**

**A** Tradeseq analysis of HSC marker genes for the B cell (red), erythroid (green), monocytic (blue) and plasmacytoid dendritic cell (pDC, purple) lineages. **B** Tradeseq analysis of lineage-specific marker genes. **C** Tradeseq analysis plots of isodecoders for all lineages.

#### Figure S7

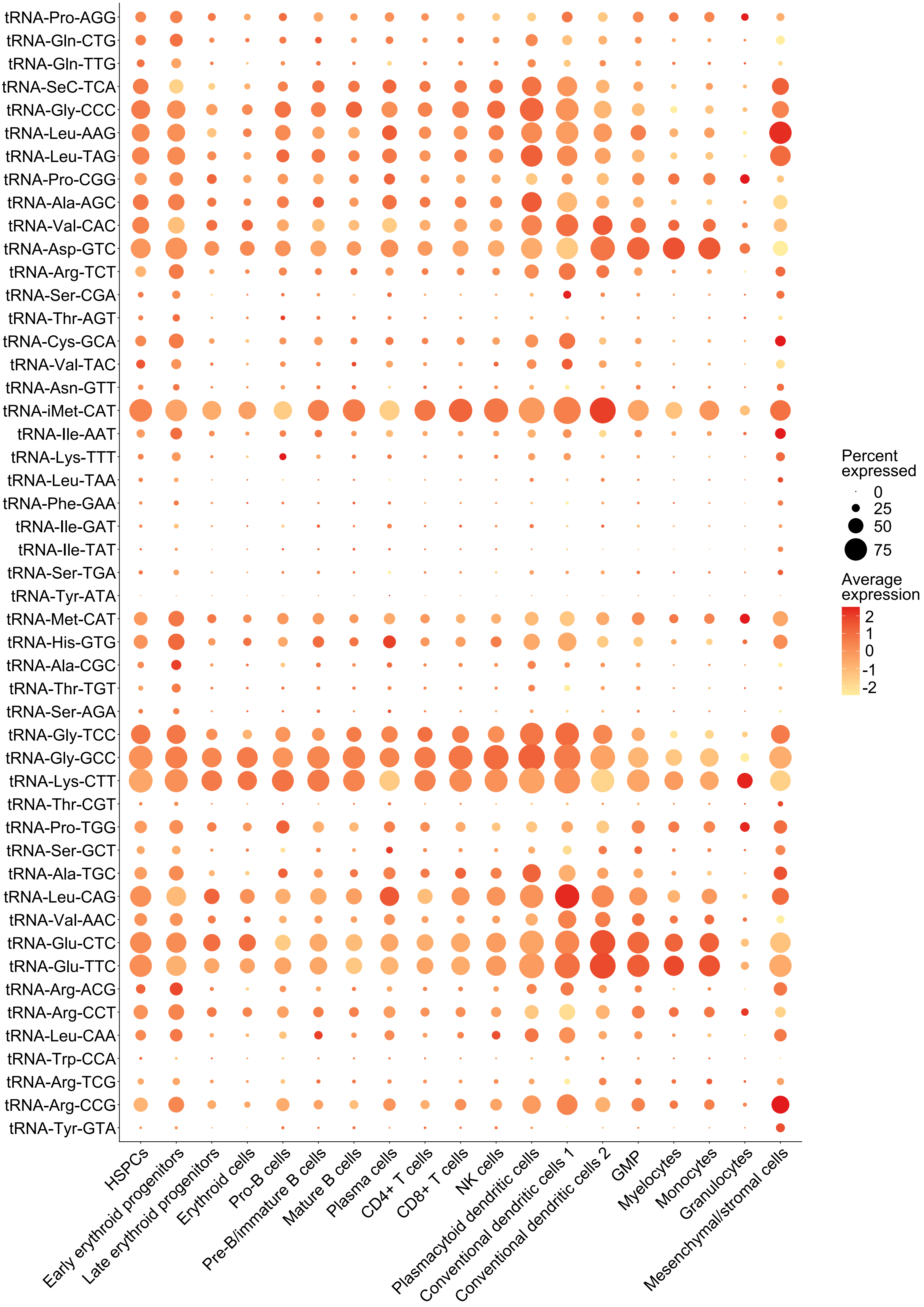

tRNA-Pro-AGG
tRNA-Gln-CTG
tRNA-Gln-TTG
tRNA-SeC-TCA
tRNA-Gly-CCC
tRNA-Leu-AAG
tRNA-Leu-TAG
tRNA-Pro-CGG
tRNA-Ala-AGC
tRNA-Val-CAC
tRNA-Asp-GTC
tRNA-Arg-TCT
tRNA-Ser-CGA
tRNA-Thr-AGT
tRNA-Cys-GCA
tRNA-Val-TAC
tRNA-Asn-GTT
tRNA-iMet-CAT
tRNA-Ile-AAT
tRNA-Lys-TTT
tRNA-Leu-TAA
tRNA-Phe-GAA
tRNA-Ile-GAT
tRNA-Ile-TAT
tRNA-Ser-TGA
tRNA-Tyr-ATA
tRNA-Met-CAT
tRNA-His-GTG
tRNA-Ala-CGC
tRNA-Thr-TGT
tRNA-Ser-AGA
tRNA-Gly-TCC
tRNA-Gly-GCC
tRNA-Lys-CTT
tRNA-Thr-CGT
tRNA-Pro-TGG
tRNA-Ser-GCT
tRNA-Ala-TGC
tRNA-Leu-CAG
tRNA-Val-AAC
tRNA-Glu-CTC
tRNA-Glu-TTC
tRNA-Arg-ACG
tRNA-Arg-CCT
tRNA-Leu-CAA
tRNA-Trp-CCA
tRNA-Arg-TCG
tRNA-Arg-CCG
tRNA-Tyr-GTA
HSPCs
Early erythroid progenitors
Late erythroid progenitors
Erythroid cells
Pro-B cells
Pre-B/immature B cells
Mature B cells
Plasma cells
CD4+ T cells
CD8+ T cells
NK cells
Plasmacytoid dendritic cells
Conventional dendritic cells 1
Conventional dendritic cells 2
GMP
Myelocytes
Monocytes
Granulocytes
Mesenchymal/stromal cells
Percent expressed
0
25
50
75
Average expression
2
1
0
-1
-2

**Fig. S7. tRNA isodecoder expression synopsis.**
A dotplot visualizing the average expression per isodecoder by color and the proportion of cells expressing the isodecoder by dot size.

**Figure S8**

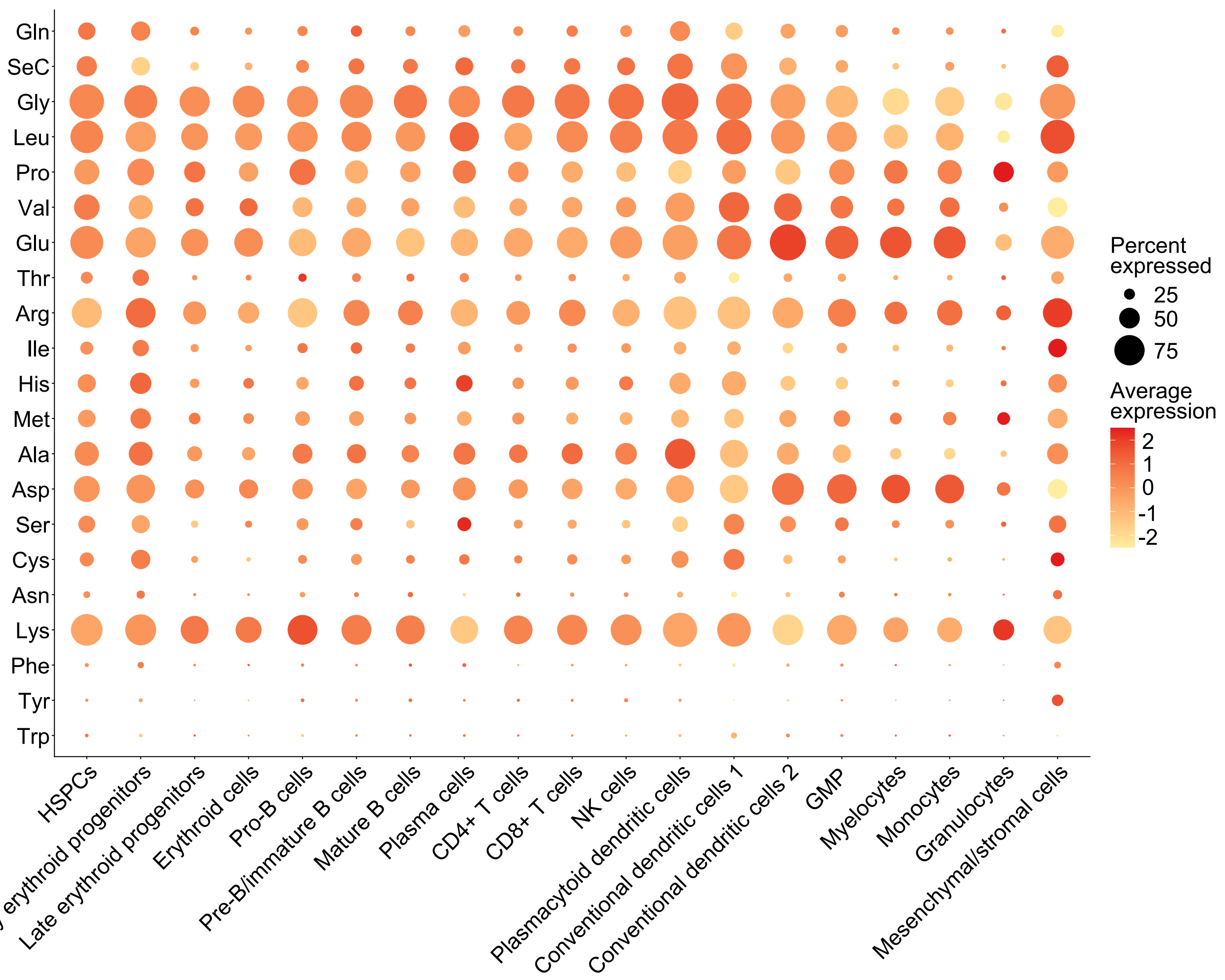

Gln
SeC
Gly
Leu
Pro
Val
Glu
Thr
Arg
Ile
His
Met
Ala
Asp
Ser
Cys
Asn
Lys
Phe
Tyr
Trp
HSPCs
Early erythroid progenitors
Late erythroid progenitors
Erythroid cells
Pro-B cells
Pre-B/immature B cells
Mature B cells
Plasma cells
CD4+ T cells
CD8+ T cells
NK cells
Plasmacytoid dendritic cells
Conventional dendritic cells 1
Conventional dendritic cells 2
GMP
Myelocytes
Monocytes
Granulocytes
Mesenchymal/stromal cells
Percent expressed
25
50
75
Average expression
2
1
0
-1
-2

**Fig. S8. tRNA isoacceptor expression synopsis.**
A dotplot visualizing the average expression per isoacceptor (amino acid) by color and the proportion of cells expressing the isoacceptor by dot size.

**Figure S9**

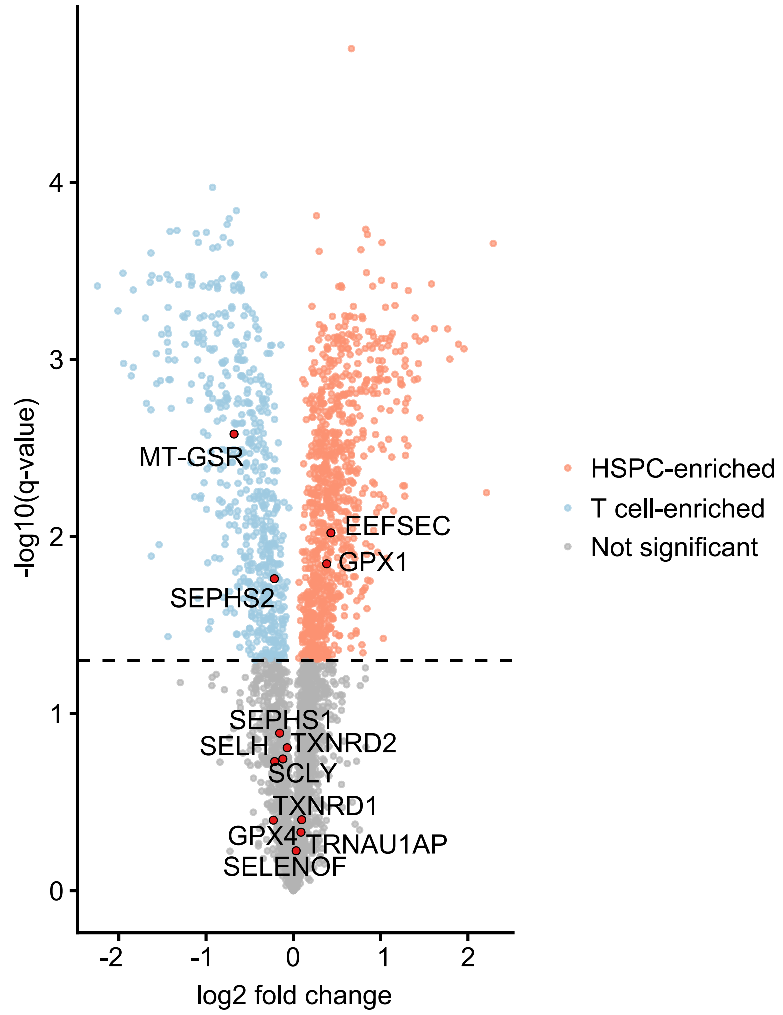
-log10(q-value)
4
3
2
1
0
-2
-1
0
1
2
log2 fold change
MT-GSR
EEFSEC
GPX1
SEPHS2
SEPHS1
SELH
TXNRD2
SCLY
TXNRD1
GPX4
TRNAU1AP
SELENOF
HSPC-enriched
T cell-enriched
Not significant

**Fig. S9. Quantitative proteomics of CD34+ HSPCs and CD4+ T cells.**
Volcano plot indicating differentially expressed selenoproteins comparing CD34+ HSPCs with CD4+ T cells from mobilized PB. padj cut-off = 0.05. Results are summarized in **supplementary file S3**.

**Figure S10**

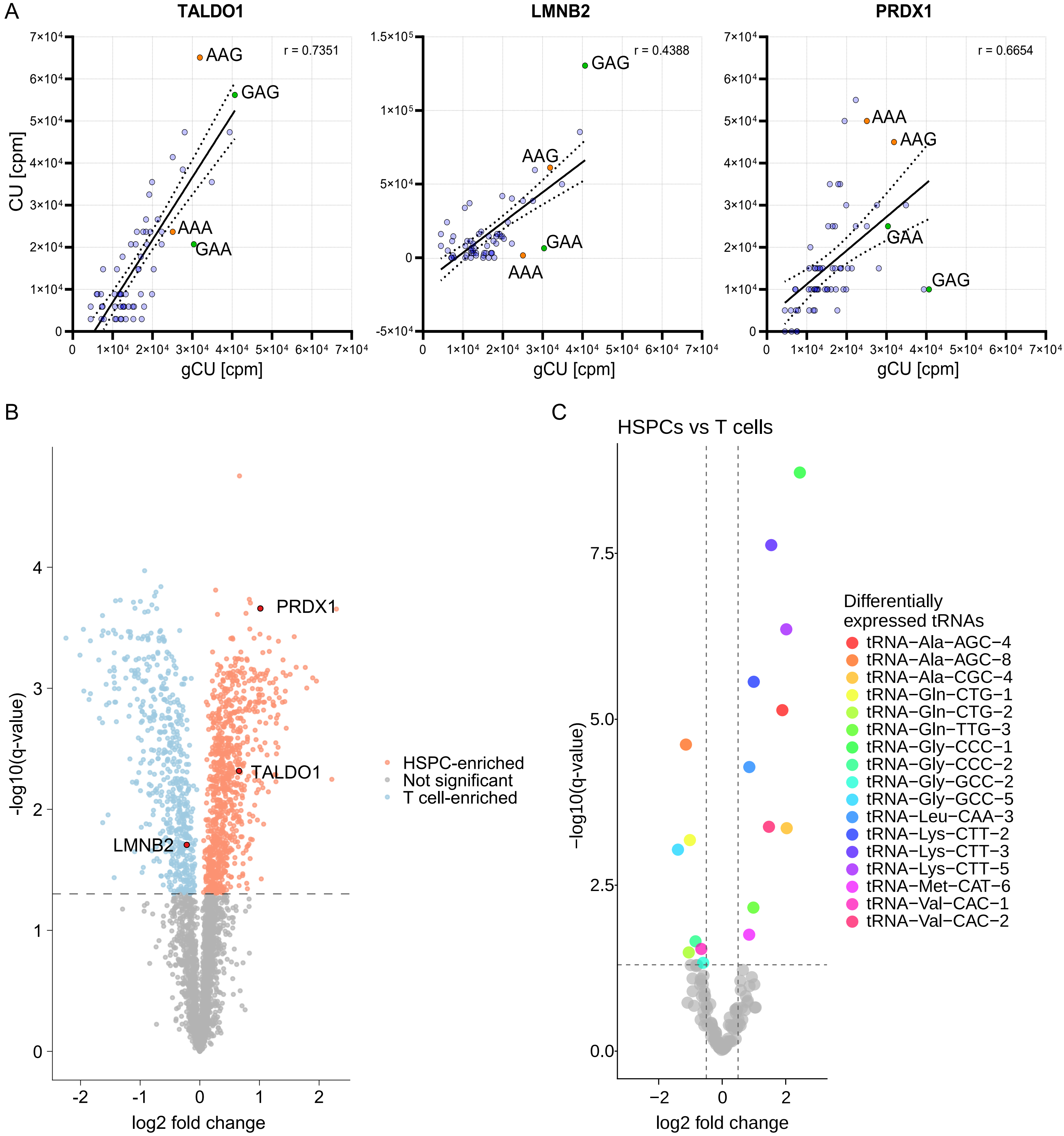
A
TALDO1
LMNB2
PRDX1
r = 0.7351
r = 0.4388
r = 0.6654
AAG
GAG
AAA
GAA
CU [cpm]
gCU [cpm]
B
PRDX1
TALDO1
LMNB2
-log10(q-value)
log2 fold change
HSPC-enriched
Not significant
T cell-enriched
C
HSPCs vs T cells
−log10(q-value)
log2 fold change
Differentially expressed tRNAs
tRNA−Ala−AGC−4
tRNA−Ala−AGC−8
tRNA−Ala−CGC−4
tRNA−Gln−CTG−1
tRNA−Gln−CTG−2
tRNA−Gln−TTG−3
tRNA−Gly−CCC−1
tRNA−Gly−CCC−2
tRNA−Gly−GCC−2
tRNA−Gly−GCC−5
tRNA−Leu−CAA−3
tRNA−Lys−CTT−2
tRNA−Lys−CTT−3
tRNA−Lys−CTT−5
tRNA−Met−CAT−6
tRNA−Val−CAC−1
tRNA−Val−CAC−2

**Fig. S10. tRNA expression and codon usage may explain quantitative protein-transcript discrepancies during HSPC differentiation.**

**A** Individual codon usage analyses of TALDO1, LMNB2, and PRDX1 open-reading-frames. Lysine codons are depicted in orange, glutamic acid codons highlighted in green. Spearman correlation factors are indicated in the top right corner of each plot. Dots indicate the mean value per codon. The line represents a simple linear regression with 95% confidence interval indicated by the dotted lines. CU codon usage. gCU genomic codon usage. cpm codons per million. **B** Volcano plot indicating differential expression of PRDX1, TALDO1, and LMNB2 comparing CD34+ HSPCs with CD3+CD4+ T cells from mobilized PB as measured by quantitative proteomics. q-value cut-off = 0.05. Results are summarized in **supplementary file S3**. **C** Volcano plot indicating differentially expressed tRNAs comparing CD34+CD38low HSPCs with CD3+CD4+ T cells as assessed using bulk tRNA-seq of FACS-isolated cells from mobilized PB. q-value cut-off = 0.05, log2fc cut-off = 0.5. Results are summarized in **supplementary file S4**.

**Figure S11**

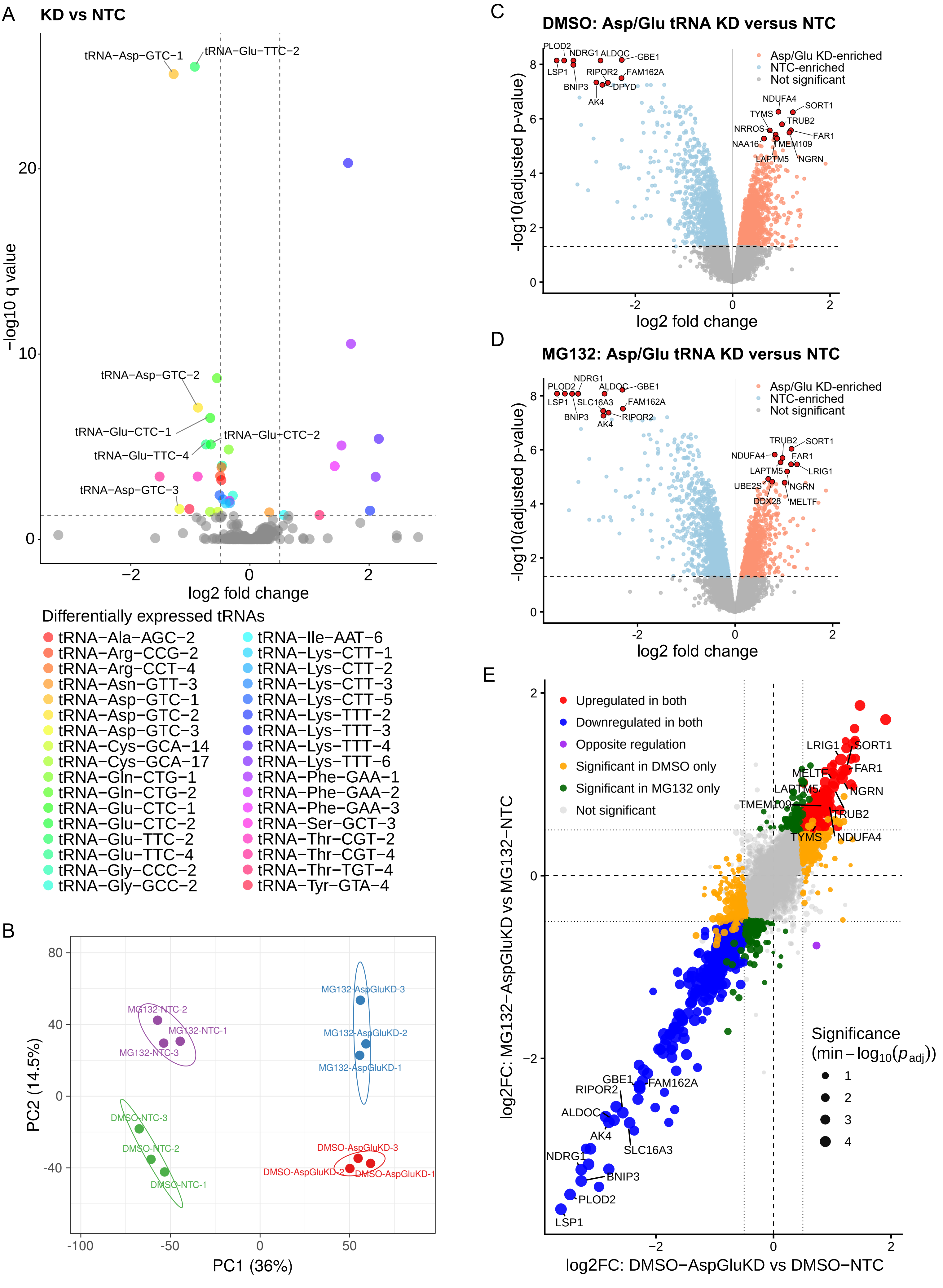
A
KD vs NTC
tRNA−Asp−GTC−1
tRNA−Glu−TTC−2
tRNA−Asp−GTC−2
tRNA−Glu−CTC−1
tRNA−Glu−CTC−2
tRNA−Glu−TTC−4
tRNA−Asp−GTC−3
−log10 q value
log2 fold change
Differentially expressed tRNAs
tRNA−Ala−AGC−2
tRNA−Arg−CCG−2
tRNA−Arg−CCT−4
tRNA−Asn−GTT−3
tRNA−Asp−GTC−1
tRNA−Asp−GTC−2
tRNA−Asp−GTC−3
tRNA−Cys−GCA−14
tRNA−Cys−GCA−17
tRNA−Gln−CTG−1
tRNA−Gln−CTG−2
tRNA−Glu−CTC−1
tRNA−Glu−CTC−2
tRNA−Glu−TTC−2
tRNA−Glu−TTC−4
tRNA−Gly−CCC−2
tRNA−Gly−GCC−2
tRNA−Ile−AAT−6
tRNA−Lys−CTT−1
tRNA−Lys−CTT−2
tRNA−Lys−CTT−3
tRNA−Lys−CTT−5
tRNA−Lys−TTT−2
tRNA−Lys−TTT−3
tRNA−Lys−TTT−4
tRNA−Lys−TTT−6
tRNA−Phe−GAA−1
tRNA−Phe−GAA−2
tRNA−Phe−GAA−3
tRNA−Ser−GCT−3
tRNA−Thr−CGT−2
tRNA−Thr−CGT−4
tRNA−Thr−TGT−4
tRNA−Tyr−GTA−4
B
MG132-NTC-2
MG132-NTC-1
MG132-NTC-3
MG132-AspGluKD-3
MG132-AspGluKD-2
MG132-AspGluKD-1
DMSO-NTC-3
DMSO-NTC-2
DMSO-NTC-1
DMSO-AspGluKD-3
DMSO-AspGluKD-2
DMSO-AspGluKD-1
PC2 (14.5%)
PC1 (36%)
C
DMSO: Asp/Glu tRNA KD versus NTC
Asp/Glu KD-enriched
NTC-enriched
Not significant
PLOD2
NDRG1
ALDOC
GBE1
LSP1
RIPOR2
FAM162A
BNIP3
DPYD
AK4
NDUFA4
SORT1
TYMS
TRUB2
NRROS
FAR1
NAA16
TMEM109
LAPTM5
NGRN
-log10(adjusted p-value)
log2 fold change
D
MG132: Asp/Glu tRNA KD versus NTC
Asp/Glu KD-enriched
NTC-enriched
Not significant
NDRG1
PLOD2
ALDOC
GBE1
LSP1
SLC16A3
FAM162A
BNIP3
RIPOR2
AK4
TRUB2
SORT1
NDUFA4
FAR1
LAPTM5
LRIG1
UBE2S
NGRN
DDX28
MELTF
-log10(adjusted p-value)
log2 fold change
E
Upregulated in both
Downregulated in both
Opposite regulation
Significant in DMSO only
Significant in MG132 only
Not significant
LRIG1
SORT1
MELTF
FAR1
LAPTM5
NGRN
TMEM109
TRUB2
TYMS
NDUFA4
GBE1
FAM162A
RIPOR2
ALDOC
AK4
SLC16A3
NDRG1
BNIP3
PLOD2
LSP1
Significance
(min−$-\log_{10}(p_{adj})$)
log2FC: MG132−AspGluKD vs MG132−NTC
log2FC: DMSO−AspGluKD vs DMSO−NTC

**Fig. S11. Reduced tRNA abundance alters the proteome.**

**A** Volcano plot of SEM cells transduced with either a tRNA Asp/Glu knockdown pool (KD) or a non-targeting control (NTC). On-target tRNAs are labeled. All quantified tRNAs are listed in **supplementary file S5**. **B** PCA based on quantitative proteomics results of Asp/Glu KD cells compared to NTC cells, treated either with proteasome inhibitor MG132 or vehicle control (DMSO). Volcano plots show differential protein expression between the tRNA Asp/Glu KD and the NTC upon DMSO (**C**) and MG132 (**D**) treatment. Raw data and differential protein expression results are summarized in **supplementary file S6**. **E** Scatterplot comparing differential protein expression between DMSO- and MG132-treated tRNA Asp/Glu KD vs. NTC. Dot size indicates the smaller -log10(padj) value of both comparisons.

#### Figure S12

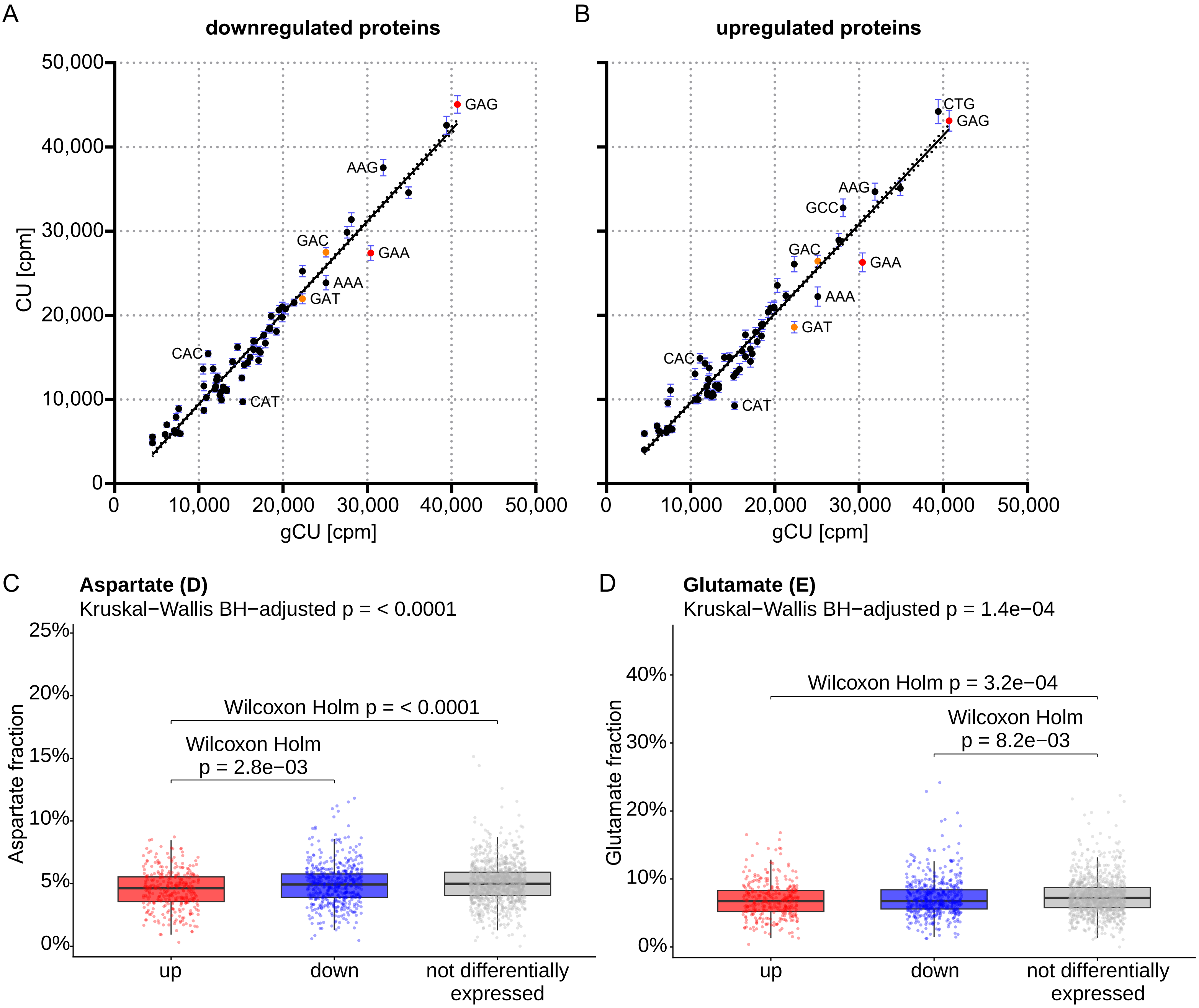

A
downregulated proteins
CU [cpm]
gCU [cpm]
GAG
AAG
GAC
GAA
AAA
GAT
CAC
CAT
B
upregulated proteins
CTG
GCC
C
Aspartate (D)
Kruskal−Wallis BH−adjusted p = < 0.0001
Wilcoxon Holm p = < 0.0001
Wilcoxon Holm
p = 2.8e−03
Aspartate fraction
up
down
not differentially expressed
D
Glutamate (E)
Kruskal−Wallis BH−adjusted p = 1.4e−04
Wilcoxon Holm p = 3.2e−04
Wilcoxon Holm
p = 8.2e−03
Glutamate fraction

**Fig. S12. Reduced tRNA availability affects protein expression in a codon- and amino acid-specific manner.**

Codon usage (CU) analysis comparing the CU of the coding sequences of all downregulated (**A**) and upregulated (**B**) proteins with the human genomic codon usage (gCU). Each dot represents the mean usage of a codon quantified as codons per million (cpm). Error bars represent the standard error of mean (S.E.M.). Asp codons are indicated in orange, Glu codons are indicated in red. The line represents a simple linear regression with 95% confidence interval indicated by the dotted lines. Amino acid composition analysis of Asp (**C**) and Glu (**D**) fractions in upregulated, downregulated, and not differentially expressed proteins. Boxplots indicate median amino acid fractions (center line) with 25th – 75th percentile interquartile ranges (box) and whiskers representing values within 1.5x the interquartile range. Each dot represents a protein. Each amino acid was tested for differences using Kruskal-Wallis corrected for multiple testing with Benjamini-Hochberg (BH) and pairwise Wilcoxon (Holm-adjusted) to identify which groups differ if the overall test result was significant. Raw data, differential protein expression results, amino acid composition analysis results incl. statistical test results are summarized in **supplementary file S6**.

**Figure S13**

A

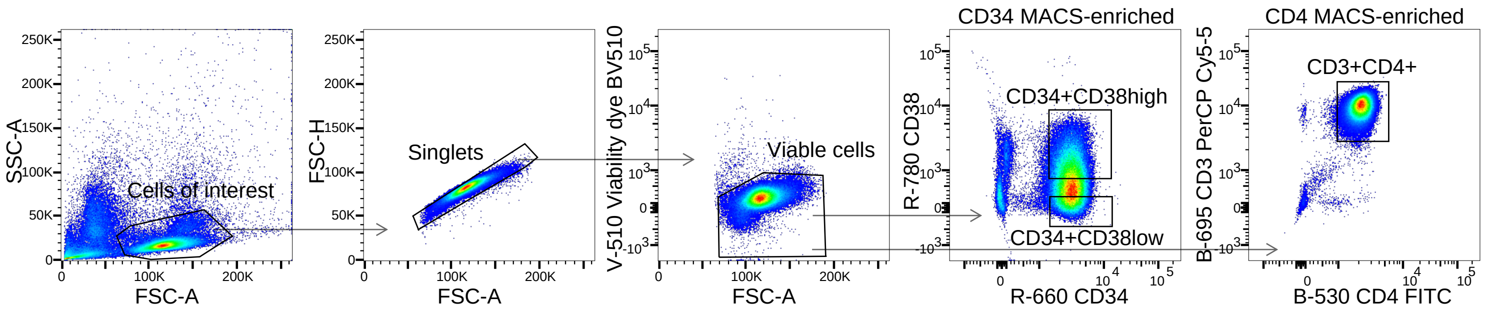

B

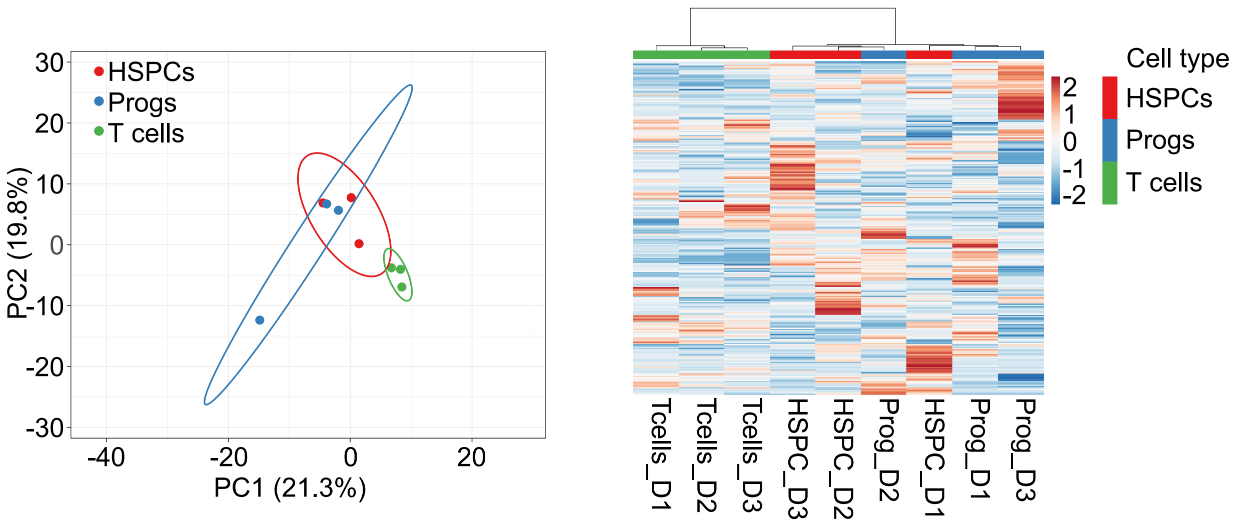

**Fig. S13. Bulk tRNA-seq of HSPCs, progenitors and T cells.**

**A** Gating strategy for the prospective FACS-isolation of CD34+CD38low HSPCs, CD34+CD38high progenitors (both from CD34 MACS-enriched PBMCs), and CD3+CD4+ T cells (from CD4 MACS-enriched PBMCs). **B** Principal component analysis (PCA) of prospectively isolated HSPCs, progenitors and T cells from three healthy donors. The prediction ellipses reflect the area in which a new sample from the same group occurs with probability 0.95. The contribution to total variation explained by each principal component (PC) is shown in percent. A heatmap visualizes the corresponding unsupervised clustering analysis.

#### Separate supplementary files

**Supplementary File S1. (separate file)**

Differential tRNA expression analyses results (DEseq2) of sc-STM-seq data in pseudobulk. Comparisons are indicated in sheet names.

**Supplementary File S2. (separate file)**

Differential tRNA expression analyses results (DEseq2) of sc-STM-seq HSPC subset data in pseudobulk. Comparisons are indicated in sheet names.

**Supplementary File S3. (separate file)**

Results of the quantitative proteomics analysis comparing CD34+ HSPCs and CD4+ T cells.

**Supplementary File S4. (separate file)**

Differential tRNA expression analyses results (DEseq2) of prospectively isolated HSPCs and T cells in bulk.

**Supplementary File S5. (separate file)**

Differential tRNA expression analyses (DEseq2) of tRNA knockdowns in comparison to a non-targeting control in bulk.

**Supplementary File S6. (separate file)**

Quantitative proteomics raw data, differential protein expression analysis, and amino acid composition analysis of the tRNA knockdowns in comparison to a non-targeting control.
